# *Escherichia coli* with a 57-codon genetic code

**DOI:** 10.1101/2025.05.02.651837

**Authors:** Wesley E. Robertson, Fabian B. H. Rehm, Martin Spinck, Raffael L. Schumann, Rongzhen Tian, Wei Liu, Yangqi Gu, Askar A. Kleefeldt, Cicely F. Day, Kim C. Liu, Yonka Christova, Jérôme F. Zürcher, Franz L. Böge, Jakob Birnbaum, Linda van Bijsterveldt, Jason W. Chin

## Abstract

The near-universal genetic code of living organisms uses 64 codons to encode the 20 canonical amino acids in protein synthesis. Here we design and generate a variant of *Escherichia coli* with a 4 Mb synthetic genome in which we replace every known occurrence of six sense codons and a stop codon with synonymous codons. We thereby recode 10^5^ codons to create an organism with a 57-codon genetic code; this organism – which we name Syn57 – uses 55 codons to encode the 20 canonical amino acids.

## Introduction

All forms of life on earth use the universal genetic code^1^, or its minimal variations^2-4^. The 64 triplet codons, in the genetic code, encode the canonical 20 amino acids and the initiation and termination of protein synthesis, and 18 of the canonical amino acids are encoded by more than one synonymous codon^5^. Synonymous codon choice is known to influence mRNA folding^6^, gene expression^7-9^, co-translational folding^10^, and protein levels^6,11,12^, and has distinct effects in distinct sequence contexts within genes and genomes^13^.

Reducing the number of codons used for genomically-encoded protein synthesis has provided a basis for generating virus resistant organisms^14-19^; it has also enabled: the more efficient synthesis of proteins containing non-canonical amino acids^14-16^, the encoded synthesis of proteins containing multiple distinct non-canonical amino acids^15,16^, and the genetically encoded synthesis of non-canonical polymers^16,20^.

Stop codons have been removed from the *E. coli* genome by mutagenesis methods^14,15^. These methods typically introduce large numbers of off target mutations^14,21^, and are not scalable to the removal of all occurrences of targeted sense codons, which are commonly orders of magnitude more abundant than stop codons^22,23^.

Emerging methods for the total synthesis of genomes^24-27^ provide opportunities to explore genome sequences that cannot be accessed by editing and are radically different from those accessed by natural evolution. To date, two bacterial genomes have been synthesized^28,29^, and efforts to synthesize other genomes^30-32^ and strategies to recode^14,21,26,30,33^, minimize^30,34^, or rearrange genomes^34-38^ are under way.

We previously reported the generation of Syn61^29^, a strain of *E. coli* with a synthetic genome in which we replaced annotated occurrences of the TCG and TCA sense codons, which normally encode serine, with defined synonyms, and the TAG stop codon with the TAA stop codon. Syn61 contains 18,214 codon changes in its genome with respect to the parental strain and uses 61 codons to make proteins. This strain has provided a foundation for creating virus-resistant cells, orthogonal genetic systems, and genetically programmed protein, polymer and macrocycle synthesis with up to three non-canonical monomers^16,18-20^. Despite the success of this strain, it remained unclear whether living organisms could tolerate much deeper sense codon compression that take them further away from natural sequence.

Here we report the total synthesis of a recoded *E. coli* genome, which implements the genome wide recoding of six sense codons and one stop codon, to deeply compress the genetic code. The resulting organism, named Syn57, has more than 10^5^ codon changes with respect to the parental strain, replaces annotated occurrences of two alanine codons and four serine codons with synonyms, and uses a 57-codon genetic code.

## Results

### Defined codon compression schemes

We empirically determined allowed defined synonymous codon recoding schemes (recoding schemes) for code compression across a 20-kb region of the *E. coli* genome rich in essential genes and target codons^24^. For each scheme we assembled the synthetic recoded DNA in a bacterial artificial chromosome (BAC), introduced it into the genome by REXER (**Supplementary Fig. 1**), and assessed the genomic recoding allowed at each position by sequencing individual post-REXER clones (**Supplementary Fig. 1a, b**).

We first tested recoding schemes in which we replaced two sense codons from the serine codon box (vS1, vS2, and vS3), or two sense codons from the alanine codon box (vA7 and vA8), with defined synonyms across the 20-kb test region (**Fig. 1a, b**). In these experiments, and all subsequent experiments, we also replaced the TAG stop codon with TAA; these experiments aimed to test genetic codes that are compressed to use three fewer codons (three-codon compression schemes). We were able to recode the 20-kb region with 2 serine schemes (vS2, vS3) and the alanine vA7 scheme, and we define these schemes as allowed recoding schemes on this region.

**Fig. 1.**
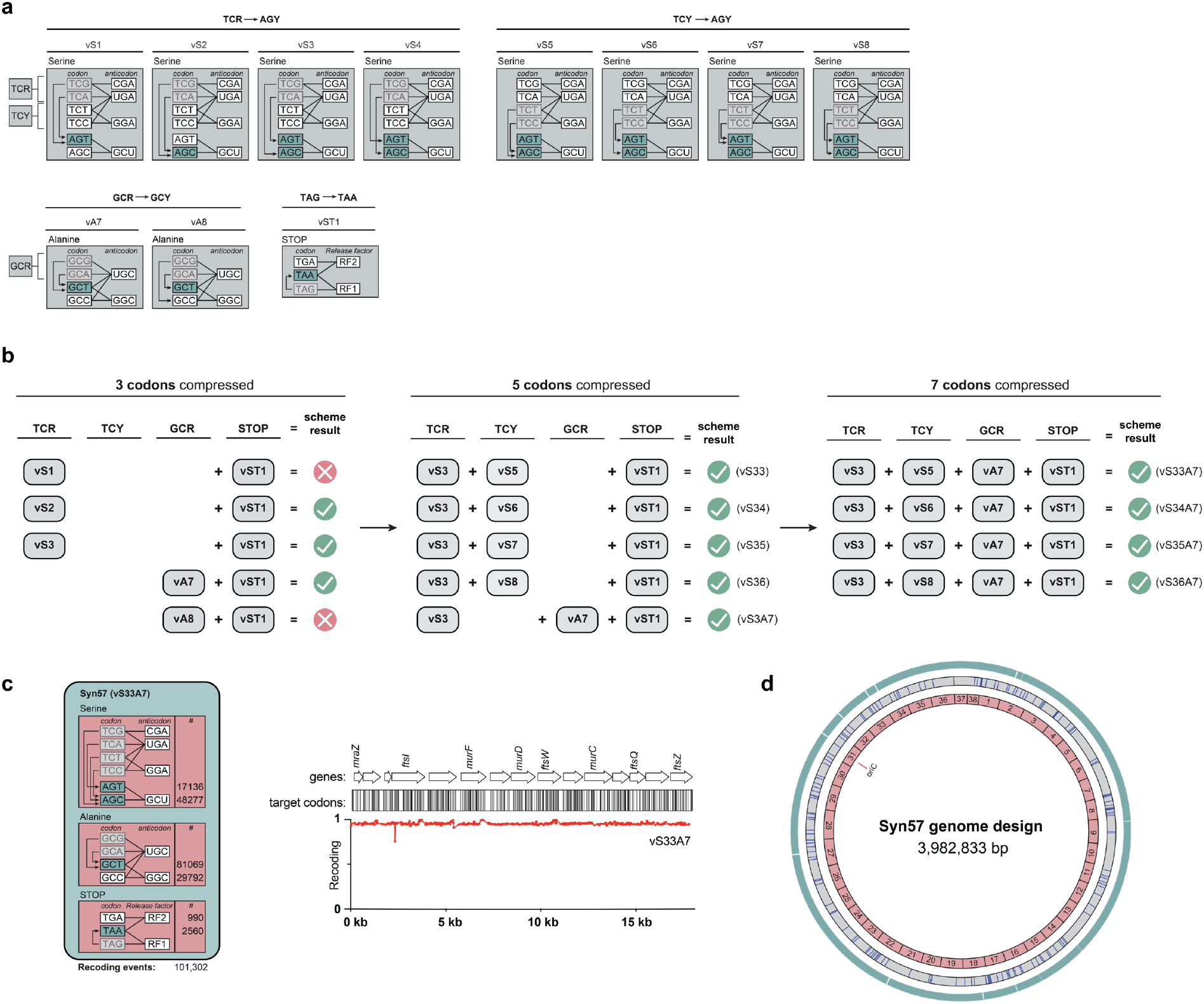
Combining codon compression schemes to yield a 57-codon genome design. **a**. Target codons to remove from the genetic code (grey boxes) were chosen from the serine and alanine sense codon boxes and the stop codon box. The recoding scheme determines which synonym (teal boxes) will replace the target codon. As input recoding schemes, we focused on the TCR and TCY codon boxes from serine, the GCR codon box from alanine, and the STOP codon box. We focused on building 3, 5 and 7 codon compression schemes by combining the indicated schemes. **b**. For 3-codon compression schemes, we combined schemes targeting reassignment of the TCR, and GCR codons with the vST1 STOP scheme. The schemes that combine vS1, vS2, vS3, and vA7 with vST1 were previously tested using REXER^24^; we additionally performed REXER with vA8 + vST1. To further compress the code by 5 codons, we combined 3-codon compression schemes targeting the TCR serine box with schemes targeting the TCY serine box (TCR + TCY) or the GCR alanine box (TCR + GCR) – this generated five different 5-codon compression schemes (vS33-vS36, and vS3A7). To remove 7 codons from the genetic code, we combined successful compression schemes from the TCR, TCY, GCR, and STOP codon boxes. Specifically, we combined the four successful TCR + TCY schemes with the GCR recoding scheme and the STOP recoding scheme – this generated four individual 7-codon compression schemes (vS33A7-vS36A7), all recoding 4 serine codons, 2 alanine codons, and 1 stop codon to different combinations of synonyms. We performed REXER with each of the indicated 3, 5 and 7 codon compression schemes (**Supplementary Fig. 2**). REXERs that led to recoding are indicated with a green tick, those that did not lead to recoding are indicated with a red X. **c**. Implementing one of the 7-codon compression schemes, vS33A7, across the entire 4-megabase (Mb) genome sequence of *E. coli* MDS42 generates a recoded genome design with a 57-codon genetic code. The recoding landscape following REXER with the vS33A7 scheme shows full recoding across the 20-kb *dcw* cluster. **d**. Map of the synthetic 57-codon genome design with all TCG, TCA, TCC, TCT, GCA, GCG, and TAG codons removed. The outer ring (101,302 teal bars, outer wheel) indicates the position of all recoding events, while the blue bars (174) indicate refactoring events (grey middle wheel). The synthetic genome design was disconnected into 38 fragments of 100 kb each (pink inner wheel).

Next, we created five new recoding schemes. In these schemes, two serine codons and two alanine codons were recoded (vS3A7, combining recoding schemes that were allowed individually), or four serine codons were recoded (vS33-vS36). We were able to recode the 20-kb region with all five schemes (**Fig. 1b, Supplementary Fig. 2a**), thereby demonstrating five allowed defined recoding schemes, each of which compresses the genetic code used on this region to use five fewer codons (5-codon compression schemes).

Finally, we combined each of the serine recoding schemes from the 5-codon compression schemes (vS33-vS36) with the alanine recoding scheme (vA7) to generate four distinct 7-codon compression schemes (**Fig. 1b, c, and Supplementary Fig. 2b**). We were able to recode the 20-kb region tested with all 7-codon compression schemes (**Fig. 1c, Supplementary Fig. 2b, and Supplementary Data 1**). We chose the vS33A7 scheme for all further work.

### Design of a codon compressed genome

We designed a 4 Mb *E. coli* genome using the vS33A7 recoding scheme. Our design recoded 101,302 target codons to defined synonyms throughout the open reading frames of the genome (**Fig. 1c, 1d**). Our genome design also included 174 refactoring events that duplicated overlapping genes containing target codons to enable their independent recoding (**Fig. 1d**). We performed a retrosynthesis on the designed genome^29^, by dividing the designed genome into 38 fragments of ∼100 kb each, and dividing each fragment into 9-14 stretches of ∼10 kb each.

### Testing defined code compression genome-wide

We obtained 10-kb stretches of synthetic DNA (from a commercial vendor) that implement our design and cover the genome. We then used homologous recombination in *S. cerevisiae* to assemble BACs containing each ∼100 kb synthetic fragment from the corresponding synthetic stretches.

We investigated the replacement of each 100-kb fragment of the genome with the corresponding synthetic fragment in 38 parallel strains. We did this using uREXER, a variant of REXER^24^ employing universal CRISPR/Cas9 spacers^25^ to excise the synthetic DNA from the BAC, and lambda-red-mediated recombination to facilitate its genomic integration in a *ΔrecA* background (**Supplementary Fig. 1c, d**).

27 of the 38 100-kb fragments led to complete replacement of the corresponding 100-kb fragment of the genome with synthetic DNA (**Fig. 2a**). These 27 fragments comprise 2.96 Mb of the entire 3.98 Mb genome and 72,826 of 101,553 total recoding events (**Fig. 2b**).

**Fig. 2.**
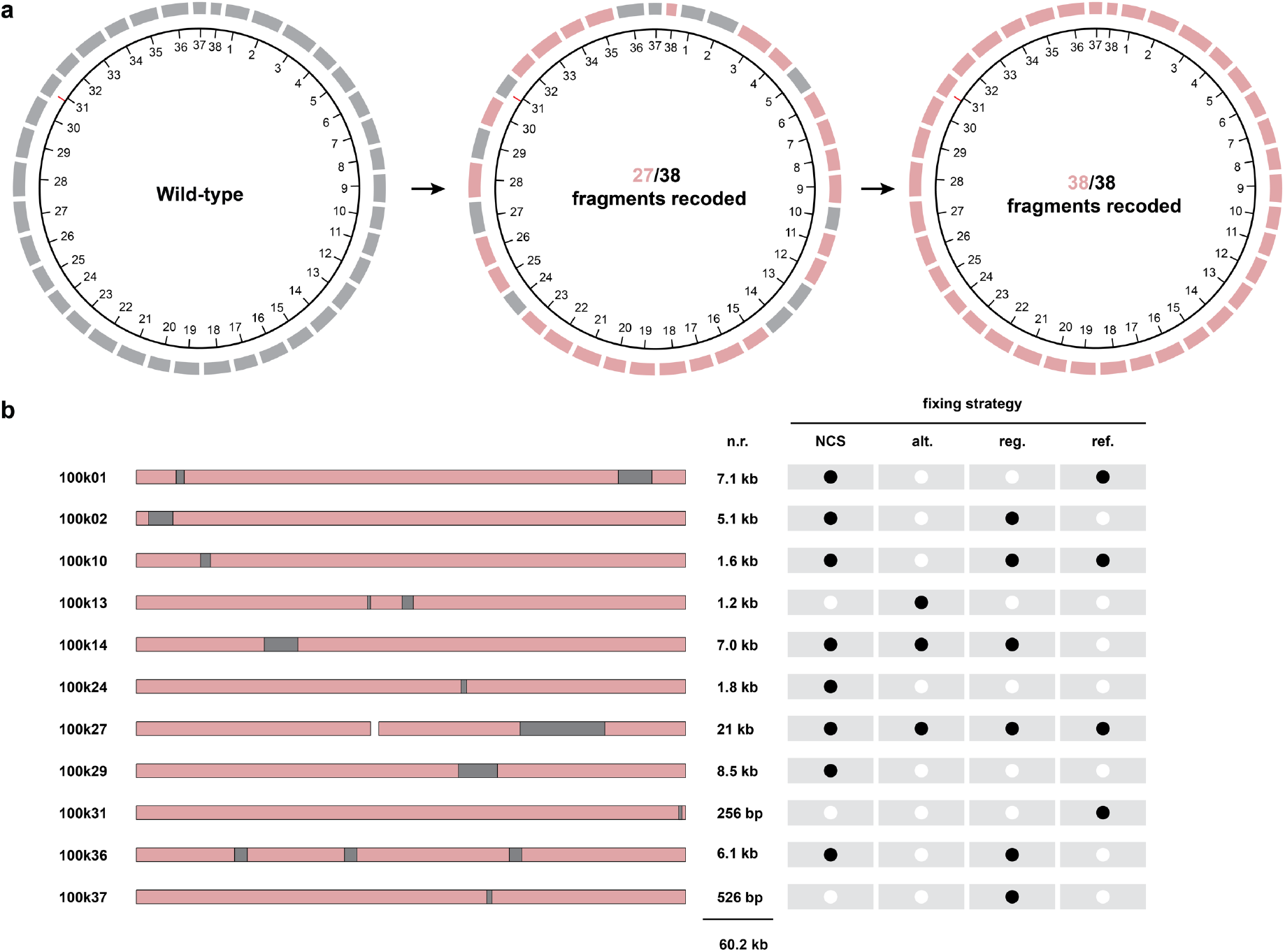
Recoding every 100-kb fragment in the genome. **a**. The genome design was split into 38 fragments, each fragment spans approximately 100 kb. The corresponding fragments on the parent genome (left) are shown in grey. Parallel uREXER experiments in wildtype cells were performed for each fragment with BACs containing synthetic DNA inserts that implement the corresponding genome design. 27 fragments (pink, middle) were fully recoded, while 11 fragments (grey) were either not recoded (100k27) by uREXER or only recoded across a part of their sequence. The ten partially recoded fragments were then further examined individually and the regions that were not replaced with the original synthetic genome design were identified, refined, and recoded according to one of four fixing strategies. This led to the recoding of all 38 fragments (right), with fragment 27 recoded across two sub-fragments (27A and 27B). **b**. For each of the 11 100-kb fragments with refined regions, the sequence that was recoded with synthetic DNA is shown in pink, and the region that remained non-recoded is shown in grey. The total length of non-recoded (n.r.) sequence within each fragment is indicated. Non-recoded regions were recoded using the fixing strategies indicated by a black dot: N-terminal coding sequence libraries (NCS), application of an alternative recoding scheme (alt.), rational fixing of regulatory modules (reg.) (e.g. promoters), or adjusting the refactoring length for previously overlapping genes (ref.). 100k27 was split into the sub-fragments 27A (left pink bar) and 27B (right pink and grey bar).

Regions of synthetic DNA that do not support cell growth as the sole copy in the genome are crossed out in post-uREXER clones; this generates chimeras between synthetic DNA and wildtype DNA. Compiled recoding landscapes^24,25,39^, generated from the sequencing of post-uREXER clones, allow us to localize the regions that maintain wildtype sequence (non-recoded) in all clones; these regions encompass regions where the synthetic sequence tested is disallowed. Within ten of the eleven remaining fragments for which we did not directly identify fully recoded clones by uREXER (01, 02, 10, 13, 14, 24, 29, 31, 36, and 37), the compiled recoding landscapes allowed us to identify regions in the genome where wildtype DNA was maintained (**Supplementary Figs. 3 to 12**).

For fragments 02, 10, 13, 14, 24 we further refined the region where the originally designed synthetic sequence is disallowed. We achieved this by performing uREXERs on the wildtype genome using (mini)BACs that contained recoded synthetic sequence covering the region of non-recoded sequence identified in the compiled recoding landscapes from the original 100-kb uREXERs. These experiments generated higher resolution compiled recoding landscapes covering each region and narrowed the regions where non-recoded DNA was maintained and where the original design for synthetic, recoded, DNA was disallowed (**Supplementary Figs. 4 to 8**).

For fragment 27 (100k27) we did not obtain any correctly phenotyping clones following uREXER (**Supplementary Fig. 13**). We mapped and compiled a recoding landscape from these experiments which clearly defined a 21 kb region containing ribosomal operons, in the 3’ half of 100k27, that was never recoded. We split the 100k27 sequence across two uREXER BACs: 27A contains 61 kb of synthetic sequence from the 5’ half of 100k27 and, 27B contains 64 kb of synthetic sequence from the 3’ half of 100k27. uREXER experiments with the 27A BAC led to complete recoding of the corresponding genomic region (**Supplementary Fig. 14**).

Overall, we recoded 3.92 Mb of the genome across 39 strains. This left 60 kb of the genome, across the 11 fragments, that was recalcitrant to replacement with the original synthetic genome design (**Fig. 2b**).

### Recoding the whole genome in 100-kb fragments

Next, we focused on discovering allowed synthetic sequences, which are consistent with our codon compression scheme, for the regions that were recalcitrant to replacement with the original synthetic genome design. We used a number of approaches, individually or in concert, to address this challenge within the eleven fragments where these regions occur; these approaches included: selecting alternative N-terminal coding sequences (NCS), using alternative recoding schemes which are consistent with code compression, altering promoter and regulatory sequences potentially disrupted by recoding, and adjusting initial refactoring designs (**Fig. 2b, Supplementary Figs. 3 to 12, 15**)^40^.

N-terminal coding sequences are important for gene expression^6,13,41-44^. We hypothesized that using alternative N-terminal coding sequences (encoded in the first eight to ten codons of genes), while maintaining recoding and code compression, might favor recoding of otherwise non-recoded sequences. We generated synthetic DNA libraries which varied the N-terminal regions of all genes, or a rationally chosen subset of genes (e.g. essential genes) within previously non-recoded regions. We aimed to replace wildtype genomic sequence with the corresponding library of variants. The NCS libraries were recombined into the (commonly wildtype) genome to identify NCS sequences that enabled full recoding across previously non-recoded regions. This strategy was used to recode the previously non-recoded region in the genomic fragment corresponding to fragment 24 (**Supplementary Fig. 8**), and contributed to the full recoding of the genomic fragments 01, 02, 10, 14, 27B, 29, and 36 (**Supplementary Figs. 3, 4, 5, 7, 15, 9, 11**).

The recoding scheme in our genome design, vS33A7, is one of several schemes that replace the same 7 target codons with synonyms, and several alternative synonymous codon compression schemes that replace the same seven codons were viable in the original 20-kb test region (**Supplementary Fig. 2b**). We used an alternative recoding scheme (vS34A7) to recode the region of the genome in fragment 13 that was not recoded with the vS33A7 scheme used in the original design. Alternative recoding schemes were also used as part of strategies that led to the full recoding of fragments 14 and 27B in the genome (**Supplementary Figs. 7, 15**).

In some cases, recoding events within annotated open reading frames overlap with the sequences of annotated promoters or other sequences that regulate gene expression. Within fragments 02, 10, 27B, and 37, such regulatory sequences are found within the regions that cannot be recoded using the original genome design. These regulatory sequences were targeted for mutagenesis as part of strategies that led to the full recoding, with code compression, of fragments 02, 10, 27B, and 37 in the genome (**Supplementary Figs. 4, 5, 15, 12**).

In four cases (located in fragments 01, 10, 27B and 31) the refactoring in our original genome design was not sufficient to enable recoding. By either extending or omitting the refactoring, we were able to fully recode these regions of the genome that could not be recoded by the original design (**Fig. 2b, Supplementary Figs. 3, 5, 10, 15**).

We found that the initial strain containing fragment 31 was not sufficiently fit for subsequent steps (**Fig. 3**). Mapping experiments demonstrated that recoding of a 4.2 kb region encompassing the ATP operon was associated with slow growth (**Supplementary Figs. 10, 16**). The fitness of this strain was improved, using NCS libraries covering the ATP operon and altered regulatory sequences (**Fig. 3b, Supplementary Figs. 17, 18**). The fitness improved strain was used for further conjugation-based assemblies.

**Fig. 3.**
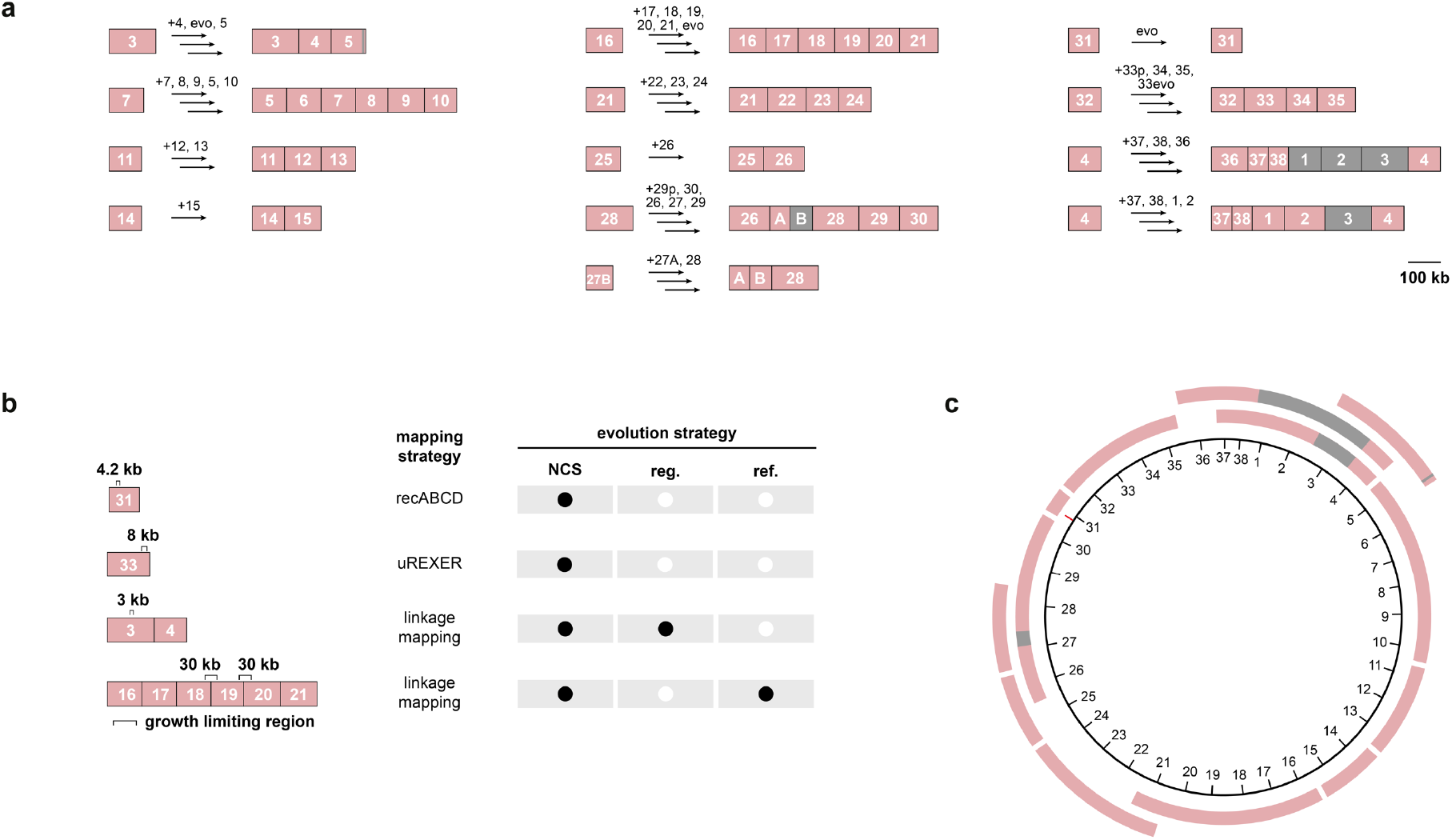
Assembly and fitness improvement of larger recoded genomic sections. **a**. The GENESIS (iterative uREXER) steps used to assemble larger recoded sections and the directed evolution steps used to improve fitness. The fragment number used in each step and directed evolution (evo) steps are indicated by a black arrow starting from individually recoded sections. Recoded sections are depicted in pink, non-recoded sections are depicted in grey, an apostrophe indicates evolved sections. The scalebar indicates 100 kb of genomic DNA. **b**. We evolved four strains with recoded fragments or sections before their use in conjugative assembly. We utilized three different mapping strategies to define growth-limiting recoded regions as targets for evolution (**Supplementary Figs. 10, 19, 21, 22**). The location and size (in kb) of the growth-limiting region is shown as a black bracket. The growth-limiting regions were targeted using the evolution strategies indicated by a black dot: N-terminal coding sequence libraries (NCS), rational fixing of regulatory modules (reg.) (e.g. promoters), or adjusting the refactoring length for previously overlapping genes (ref.) (**Supplementary Figs. 18, 19, 21, 22**). **c**. The recoded sections in the thirteen strains generated by GENESIS tiles the Syn57 genome as shown by the pink bars, tick marks show the genome fragment number.

Overall, we recoded the entirety of each 100-kb fragment (**Fig. 2a**), such that the sequence of the entire genome was recoded across 39 strains.

### Synthesis of recoded sections

We used a version of genome stepwise interchange synthesis (GENESIS, iterations of uREXER) to compile recoded fragments into single strains (**Fig. 3a**). This generated 13 recoded strains, containing larger recoded genomic sections, that cover the genome (**Fig. 3a, c**). In the process of generating these recoded sections we fixed 100k33 (**Supplementary Fig. 19**) to enable complete recoding of 100k32-35 (r32-35) and, improved the growth of r03-04 and r16-21, as described below.

### Fixing and enhancing recoded sections from GENESIS

The compiled recoding landscape from post-uREXER clones for adding 100k33 to 100k32 in the genome identified an 8-kb region within 100k33 that was not recoded in the resulting genome (**Supplementary Fig. 19a**); NCS fixes within this region enabled complete genome recoding within r32-35 (**Fig. 3b, Supplementary Fig. 19**).

The complete recoding of 100k03 and 100k04 (r03-04) in a single genome and the complete recoding of r16-21 led to slow growing cells (**Supplementary Fig. 16**). We therefore wanted to identify regions of recoded sequence that limit growth of these cells, and use this information to target these regions for fixing.

To pinpoint sequences within larger recoded sections that are associated with fitness defects, we generated linkage maps^45,46^ between recoding and fitness. We used bacterial conjugation to transfer a genome section from a parental, non-recoded (donor) strain into a (recipient) strain harbouring the recoded section of interest. After conjugative transfer of the wildtype DNA, RecABCD recombination generated genomes which contain chimeras between wildtype sequence and recoded sequence. These chimeras may have varying degrees of recoding and varying degrees of fitness (**Supplementary Fig. 20**).

We then subjected the strains with chimeric genomes to growth selection, to enrich faster growing cells. At regular intervals we isolated individual clones and measured their growth curves and determined the extent of their recoding by NGS. We collected this data for numerous clones with diverse fitness.

The data from all clones was compiled into a recoding-fitness linkage map (**Supplementary Fig. 20**). For each recoded codon position in the recipient genome we plotted the limit value for a fitness metric when recoding is maintained at that position in chimeric genomes; for example, we plotted the lowest doubling time for all clones that maintain recoding at each position. This reveals the location and severity of defects within the recoded sequence, and has the potential to provide resolution down to single recoded codons.

For the strain with r03-04 recoded we compiled sequencing and growth data from 88 unique clones into a recoding-fitness linkage map. The linkage map revealed a 3-kb region within which recoded positions limited growth. NCS and regulatory fixes within this region substantially enhanced growth (**Supplementary Figs. 16, 21**).

In the case of the strain r16-21, we compiled sequencing and growth data from 128 unique clones into a recoding-fitness linkage map. The linkage map highlighted two genomic regions, of about 30 kb each, centred on the borders between fragments 18 and 19, and between fragments 19 and 20 (**Supplementary Fig. 22**). Recoding in these regions was linked to slower growth. This linkage map therefore pinpointed two distinct regions where recoding limited growth (**Supplementary Fig. 22**). The structure of the growth associated *nuo* operon, within the growth limiting region between 18 and 19, suggests that it utilizes the termination-reinitiation mechanism for operon translation. We predicted that a refactoring event in our genome design, which inserted 20 bases between *nuoJ* and *nuoK*, might disrupt translation of genes downstream of *nuoJ*; we therefore altered this refactoring event to a sequence predicted to re-instate the desired termination-reinitiation. Within the growth limiting region between 19 and 20 we identified two essential genes (*ligA* and *zipA*) and targeted these for NCS fixes. The NCS fixes and refactoring alterations within these two regions enhanced growth (**Supplementary Figs. 16, 22**).

The fixed and improved growth strains with r32-35, r03-04, and r16-21 recoded, along with the other strains generated by GENESIS (**Fig. 3c**), were used for conjugative assembly of larger recoded genomic sections.

### Assembly of a recoded genome

We used conjugative transfer and recombination (conjugative assembly) to recombine recoded genome sections, generated by GENESIS, from donor strains into adjacent recoded genome sections in recipient strains^29,47^(**Supplementary Fig. 23**). Origin of transfer (*oriT*) cassettes were installed at one end of the recoded sequence in donor strains, such that they transferred the recoded sequence to the recipient stain. To ensure efficient recombination, recipient cells contained at least 3 kb of recoded sequence that overlapped with the recoded sequence adjacent to *oriT* in the donor cells; this sequence was followed by a double selection cassette. We selected for recipient cells which had lost the negative marker at the end of the recoded recipient sequence and gained the positive marker at the end of the donated recoded sequence.

We used a convergent series of conjugative assembly steps to compile the recoded sequence into a single genome (**Fig. 4**). At several points in the series we mapped and/or evolved the strains to generate conjugative donors or recipients that were competent for the next assembly step (**Fig. 4, Supplementary Figs. 24 to 28**). For example, the evolution of r03-31(-27B) (**Fig. 28**) was crucial for enabling the conjugation to generate a robust Syn57(-27B) strain; this strain was then conjugated with r24-30, which was derived from r24-30(-27B) via uREXER of a 27B sequence (**Supplementary Fig. 29**), to generate the final strain.

**Fig. 4.**
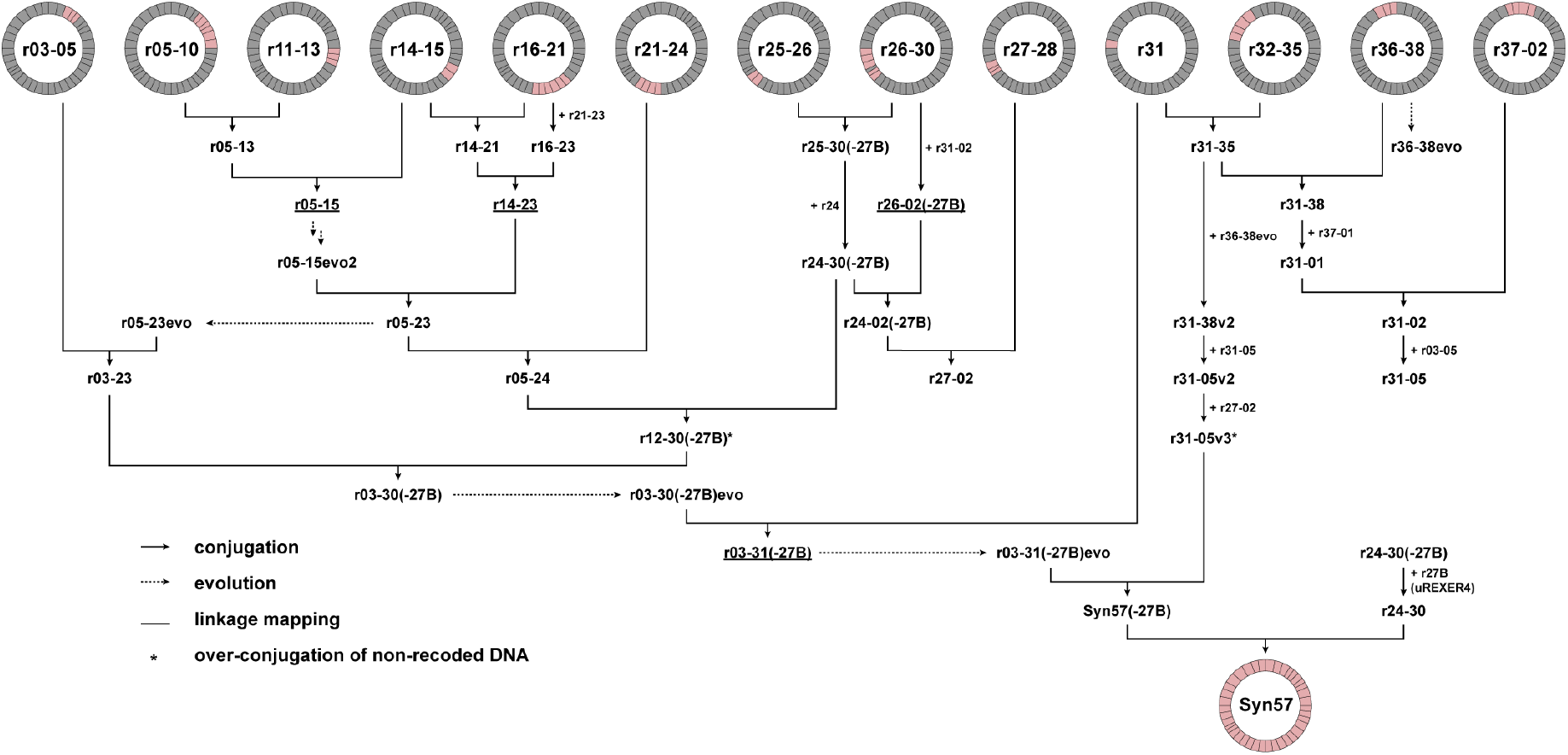
Total genome assembly to create Syn57. Starting with the set of 13 partially synthetic genome strains (shown in **Fig. 3**), we assembled a 57-codon genome into a single fully recoded strain using the indicated sequence of conjugations (black arrows). Donor and recipient strains contain unique recoded genomic sections (denoted in pink), with the donor stain harbouring a recoded genomic section downstream of the recoded genomic section in the recipient strain. Conjugation proceeded with interspersed steps of linkage mapping (underlined sections) and evolution (dotted arrows) to optimize strain fitness prior to downstream genome assembly steps, which ultimately yielded the final fully recoded strain Syn57. Over-conjugation during efforts to generate r27-05, via conjugation of a r31-05v2 donor on a r27-02 recipient, yielded r31-05v3 containing a duplication within r33. We then utilized r31-05v3 to generate Syn57(-27B) in preparation for the final conjugation step to transfer r27B.

We named the final synthetic *E. coli* ‘Syn57’. In this strain all 1.01 × 10^5^ target codons from the parental genome are recoded. 107 target codons are recoded to synonyms consistent with code compression but, distinct from those in the original genome design and, 4 of the 174 refactoring events in the original design are deleted in Syn57. Thus, 99.9% of the target codons were recoded according to the original design and 98% of the refactoring events are maintained from the original design. Syn57 contains 307 mutations, and 15 indels at non target codons; the vast majority of these mutations were programmed into strain assembly intermediates.

## Discussion

We have demonstrated the high-fidelity total synthesis of a functional genome for an organism with a 57-codon genetic code. The synthetic genome contains more than 10^5^ codon changes with respect to the parental genome, and replaces annotated occurrences of seven codons – four serine codons, two alanine codons, and the amber stop codon with synonyms for code compression. TCN codons for serine were converted to AGT/C codons and GCR codons for alanine were converted to GCT/C codons; these changes converted both the six-codon box for serine and the four-codon box for alanine to two-codon boxes. This work exemplifies how genome synthesis can move the genome sequences of organisms into new regions of sequence space that may not have been accessed by natural life.

The synthesis of Syn57 proceeded through the recoding of 100-kb fragments (that cover the whole genome) across 39 strains by uREXER, the assembly of these fragments into sections by GENESIS, the conjugative assembly of these sections into larger sections and, ultimately the completion of the recoded genome. At each stage of the synthesis we developed and implemented strategies to map the locations of defects in the original design. Defects within 100-kb synthetic fragments were mapped using uREXER and RecABCD mediated recombination, while defects arising in larger sections were identified using variants of conjugative linkage mapping. These mapping experiments provided the targets for fixing the synthetic sequences, and the mapping and fixing at each stage of the synthesis was often crucial to enabling the next step of the synthesis. These experiments provide a paradigm for integrating ‘just in time’^48^ defect mapping and fixing of initial designs into synthetic schemes, such that local defects are identified and fixed early in the synthesis and longer range, potentially epistatic or synthetic lethal, defects are identified and fixed as they emerge in the assembly process.

In future work we will build on the generation of the deeply recoded strain we have created to explore the generation of deeply orthogonal genetic codes, enhanced virus resistance, the genetic encoding of up to seven distinct non-canonical monomers into proteins and the encoded cellular synthesis of non-canonical macrocycles and polymers composed of up to seven new monomers.

## Acknowledgments

We thank Steven Wingett for assistance with Nextflow. We thank Martyn Howard and Mark Cussens from the LMB Media Kitchen.

## Author contributions

W.E.R. tested recoding schemes and generated the initial genome design. W.E.R., F.B.H.R., M.S., R.L.S., R.T., W.L., Y.G., A.A.K., K.C.L., J.F.Z., J.B., L.vB. assembled BACs and/or performed uREXER experiments. R.T. directed the use of N-terminal coding sequence libraries. W.E.R., F.B.H.R., M.S., R.L.S., R.T., W.L., Y.G. fixed non-recoded regions in individual fragments. M.S. developed and directed genomic integrations and refinement of defective sections, and linkage mapping for longer recoded sections. M.S., W.L., A.A.K performed mapping experiments. R.T. recoded the 27B fragment. W.E.R., R.L.S., C.F.D., K.C.L. coordinated RNA-Seq experiments. F.B.H.R. developed and directed oligo-based evolution for fitness improvement. F.B.H.R., M.S., A.A.K., C.F.D. performed oligo-based evolutions. W.E.R., F.B.H.R., M.S., R.L.S., R.T., C.F.D. performed conjugative assemblies of sections. R.L.S., Y.G., C.F.D., K.C.L., F.L.B. wrote and/or refined scripts for data analysis. F.B.H.R., M.S., R.L.S., A.A.K., Y.C. performed growth measurements. J.W.C. supervised the project. J.W.C. wrote the manuscript with input from W.E.R., F.B.H.R., M.S., R.L.S., R.T.

## Supplementary Materials

Materials and Methods

Supplementary Figures 1-29

Supplementary Data 1

References (*49-63*)

