## Supplementary material for "*Escherichia coli* with a 57-codon genetic code": Methods and Supplementary Figures

**This PDF file includes:**

Materials and Methods
Supplementary Figs. 1 to 29
Supplementary References

**Other Supplementary Material for this manuscript includes the following:**

Supplementary Data 1

### Materials and Methods

#### *Combining codon compression schemes*

To empirically determine deep codon compression schemes, we combined codon compression schemes targeting the serine, alanine, and stop codon boxes. In our approach, additive recoding requires combining recoding schemes which target different 2-codon boxes (**Fig. 1b**). As input 3-codon compression schemes, we combined schemes targeting the TCR (serine), TCY (serine), and GCR (alanine) 2-codon boxes with the vST1 (STOP) scheme. To generate 5-codon compression schemes, we combined 3-codon compression schemes targeting the TCR serine box with schemes targeting the TCY serine box (TCR + TCY) or the GCR alanine box (TCR + GCR). Before proceeding with further codon compression, we experimentally determined the viability of individual 5-codon compression schemes by performing REXER and compiling a recoding landscape across a 20-kb region of the *E. coli* genome rich in essential genes and target codons[49]. We used MDS42$^{rpsL\text{(K43R)}}$ cells harbouring a *rpsL-kanR* double-selection cassette as a genomic landing site at the 5' end of the 20-kb region and the pKW20_CDFtet_pAraRedCas9_tracrRNA plasmid expressing the lambda red alpha/beta/gamma genes along with Cas9[49]. To assemble the synthetic DNA harbouring the recoded sequence, we performed yeast assemblies to generate the YAC/BAC using the following input pieces: i) a YAC origin/ARS alongside a BAC origin, ii) a *sacB-cat* double-selection cassette with homology to the 3' end of the 20-kb region, and iii) two 10-kb stretches of synthetic DNA encoding the recoded 20-kb region (see below in the section '***BAC assembly and delivery***' for details on yeast assembly). After transforming an assembled BAC harbouring the recoded 20-kb region, we performed REXER as described previously[49] and took 16 post-REXER clones forward for next-generation sequencing (NGS). Schemes with at least 75% of clones with a fully recoded 20-kb region were deemed as successful codon compression schemes and were taken forward for further codon compression (**Supplementary Data 1**). This process was repeated for the 7-codon compression schemes to yield an empirically validated design for a 57-codon genetic code.

***Codon compressed genome design***

We based our synthetic genome design on the sequence of the *E. coli* MDS42 genome (accession number AP012306.1), which has 3547 annotated CDSs. From this initial genome template, we then updated our annotation based upon i) identifying discrepancies between the MDS42 genome annotation and its parental MG1655 genome annotation, and ii) comparing nucleotide sequence-based CDS length from the MDS42 genome annotation to the proteomics-based CDS length from the Uniprot database[50]. In total we updated 33 CDS annotations, mostly to include N-terminal extensions that increased the length of the CDS. Our synthetic genome annotation includes the manual downgrading of 3 CDS annotations to pseudogenes (*htgA, ybbV, yzfA*) and the manual promotion of 12 pseudogenes to CDS annotations (*ydeU*, *ygaY*, *pbl*, *yghX*, *yghY*, *agaW*, *yhiK*, *yhjQ*, *rph*, *ysdC*, *glvG*, *cybC*) – these updates were based upon proteomic evidence or lack thereof [50]. To enable negative selection with *rpsL*, we mutated the genomic copy of *rpsL* to *rpsL*$^{K43R}$.

To recode our updated MDS42 genome annotation, we used a custom Python script[51] that i) identifies and recodes all target codons to defined synonyms, and ii) identifies and resolves overlapping gene sequences that contain target codons. From our curated MDS42 starting sequence, we used the script to generate a recoded synthetic genome in which all TCG, TCA, TCT, TCC, GCG, GCA and TAG codons were replaced with AGC, AGT, AGC, AGC, GCT, GCT, and TAA codons, respectively.

***Retrosynthesis of recoded stretches***

We divided the ~4 Mb designed genome into 38 fragments of between 60 and 136 kb. We designated the boundary sequences between these fragments as 'landing sites.' For each fragment, the landing site is located at its 3' end and overlaps by 50-100 bp of homologous sequence to the 5' end of the downstream, neighbouring fragment.

Each of the ~100-kb fragments were broken down into ~10 stetches of ~10 kb each. To facilitate downstream assembly by yeast, we designed each stretch to overlap with an adjacent stretch by 80-200 bp of homologous sequence. We ordered a total of 409 stretches to be synthesised and delivered in vector format (Twist Biosciences, USA), which we subsequently linearized by virtue of flanking BsaI, AvrII, SpeI, or XbaI restriction digest sites.

***Construction of E. coli strains containing double-selection cassettes at genomic landing sites***

According to our design, each region of the genome that is targeted for replacement by a synthetic fragment is flanked by a landing site. The location of genomic landing sites was chosen primarily to avoid disrupting annotated genome features such as promoters or CDSs, therefore avoiding potentially deleterious effects from double-selection cassette integration. The replacement of wild-type (WT) genomic DNA with recoded synthetic DNA requires the presence of a double-selection cassette at the 5' landing site, for which we used the following negative and positive selection markers: *rpsL* (−1, streptomycin sensitivity), *kanR* (+1, kanamycin resistance), *sacB* (−2, sucrose sensitivity), *cat* (+2, chloramphenicol resistance), *pheS*$^{T251A\_A294G}$ (*pheS**; −3, 4-Chloro-phenylalanine (4-CP) sensitivity), *hygR* (+3, hygromycin resistance), *gentR* (+4, gentamycin resistance), *tetR* (+5, tetracycline resistance), *ampR* (+6, ampicillin resistance), *nrsR* (+7, nourseothricin resistance), *aprR* (+8, apramycin resistance), and *specR* (+9, spectinomycin resistance). We primarily integrated *rpsL-kanR, pheS*-hygR*, or *sacB-cat* double-selection cassettes as the upstream genomic landing site for the region of interest (e.g. LS01 *sacB-cat* for 100k02 recoded fragment integration). As described previously in detail[52], to integrate a double-selection cassette at a given genomic landing site via lambda red-based recombination, we first transformed MDS42$^{rpsL(K43R)/\Delta recA}$ cells[53] harbouring either the pKW20_CDFtet_pAraRedCas9_tracrRNA (*tetR*) or pKW20_CDFApr_pAraRedCas9_tracrRNA (*aprR*) plasmid expressing the lambda red alpha/beta/gamma genes, induced them with 0.5% L-arabinose, and prepared them as electrocompetent. We then transformed PCR products of double-selection cassettes flanked by 50 bp homology arms to the genomic landing site of interest, and following recovery we plated cells on LB agar plates supplemented with either kanamycin (50 µg/mL) for *rpsL-kanR*, hygromycin (200 µg/mL) for *pheS*-hygR*, or chloramphenicol (20 µg/mL) for *sacB-cat* cassettes.

***BAC assembly and delivery***

We constructed Bacterial Artificial Chromosome (BAC) shuttle vectors that contained 60-136 kb of synthetic DNA and all the functional components required for uREXER. The 5' side of synthetic DNA was flanked by a region of homology to the genome (HR1), and a Cas9 cut site. The 3' side of synthetic DNA was flanked by a double-selection cassette, a region of homology to the genome (HR2), a second Cas9 cut site, and the expression cassette for two universal spacers targeting the two Cas9 cut sites flanking the HR1 and HR2 regions. The BAC also contained a negative selection marker, a BAC origin, a URA marker and a YAC origin (*CEN6* centromere fused to an autonomously replicating sequence (CEN/ARS)).

We assembled BACs by homologous recombination in *S. cerevisiae* B4741 cells. Each assembly combined i) 7-14 stretches of synthetic DNA, each 6-13 kb in length; with ii) a selection construct (see below), iii) a universal spacers construct, and iv) a BAC/YAC shuttle vector backbone[49].

Synthetic DNA stretches were either excised by digestion with BsaI, AvrII, SpeI, or XbaI restriction sites, or amplified by PCR from commercially-sourced 10 kb vectors (Twist Biosciences, USA).

Selection constructs contained a region of homology to the 3' most stretch of the fragment, a double-selection cassette (*sacB-cat* or *rpsL-kanR)*, a region of homology (HR2) to the targeted genomic locus, a negative selection marker (*rpsL, sacB or pheS*)*, and the YAC origin CEN/ARS. We amplified selection constructs by PCR from previously assembled BACs[51] corresponding to the homologous fragment in the genome.

Universal spacer constructs contain two universal spacers, targeting either even or odd BACs, and enable the conversion of BACs from circular to linear dsDNA in the presence of Cas9 in cells. We generated universal spacer construct pieces with homologies for yeast assembly by PCR from a previously generated template.

For the BAC backbone, we PCR amplified the sequence containing a BAC origin and a *URA3* marker from a previously assembled BAC[49].

For BAC assembly, we utilized the natural recombinogenic ability of *S. cerevisiae*, as described previously[52,54]. To program recombination, we designed each input piece with 60-180 bp of homology to adjacent pieces. To initiate yeast assembly, we transformed *S. cerevisiae* BY4741 spheroplasts with 30-50 fmol of the following input DNA pieces: the BAC selection construct, a BAC/YAC backbone, and each piece of synthetic DNA. Following selection on ΔURA plates, we identified yeast clones harbouring correctly assembled BACs by either i) colony PCR followed by NGS, or ii) direct NGS with yeast total nucleic acid extracts (as described below).

To extract assembled BACs from yeast, we adapted the Zymo Yeast Extract Kit for a 96-well plate format. Briefly, we inoculated single yeast colonies into 300 μL of YM4/2% glucose/ΔURA media into individual wells of 1.2 mL 96-well plates (AB-1127) and incubated them overnight at 30 °C while shaking at 700 rpm (Grant Bio PHMP-4). We then spun down overnight cultures and digested them with 3.5 μL of zymolyase mixed with 80 μL of YD digestion buffer (Zymo Research) – digestion proceeded for 1 hour at 37 °C while shaking at 500 rpm. We then isolated DNA by adding 30 μL of AMPure Beads (Beckman Coulter) followed by the manufacturer's recommended protocol for washing with 70% ethanol and eluting with 50 μL of water. For yeast extracts from each well, we assessed the fidelity of BAC assembly by next-generation sequencing (NGS) (see section '***Illumina NGS data analysis***' for details). Following NGS confirmation, we electroporated the assembled BAC yeast extract into DH10b and/or MDS42$^{rpsL(K43R)/\Delta recA}$ cells and selected for the positive selection marker on the BAC when plating. We isolated the BACs from overnight cultures using a commercial miniprep kit (Qiagen) where we mixed very gently after addition of the lysis and the neutralization buffers, and we decreased the spinning speed from ~17,900 rcf to 5,000 rcf in the column binding and washing steps to avoid shearing of the BAC. For higher yields and better transformation efficiencies, we alternatively isolated the BAC from a 20 mL overnight culture via isopropanol precipitation where we followed the steps of the miniprep kit (Qiagen) but precipitated the DNA from the supernatant with isopropanol and the obtained pellet was washed with 70% ethanol. The DNA was then resuspended in a final volume of 50-100 μL of water.

***uREXER to integrate 100-kb recoded fragments***

We generated universal REXER (uREXER) BACs by adapting the design of REXER (Replicon Excision for Enhanced Recombination) BACs by adding Cas9-based universal spacers to the BAC backbone. We PCR amplified universal spacer cassettes for odd and even BACs from episomes harbouring the relevant cassettes[53], and we placed them downstream of selection constructs in uREXER BAC designs. Utilizing universal spacers cassettes fused into the uREXER BAC construct obviates the need for: i) bespoke spacer plasmid cloning[49], and ii) supplementation of spacers in trans during synthetic DNA integration experiments - this expedited the synthesis of our recoded genome.

To initiate uREXER, we transformed MDS42$^{rpsL\text{(K43R)}/\Delta recA}$ cells containing pKW20_CDFtet_pAraRedCas9_tracrRNA[49] and a double-selection cassette at the relevant upstream genomic landing site with the relevant isolated BAC (e.g. LS01 *sacB-cat* with 100k02 BAC). Following recovery for 1 hour in 1 mL of SOB, we plated transformed cells on 2xYT agar supplemented with 2% glucose, 5 µg/ml tetracycline and antibiotic selecting for the BAC (i.e. 20 µg/ml chloramphenicol for odd BACs or 50 µg/ml kanamycin for even BACs). Following restreaking of individual colonies, we inoculated colonies into 2xYT medium (5 mL) with 5 µg/ml tetracycline, 2% glucose (w/v) and the BAC-specific antibiotic, and we grew the cultures overnight at 37 °C, 220 rpm. To initiate the expression of the lambda red components and Cas9, we then spun down the overnight cultures and resuspended them into 200 mL of 2xYT medium with 0.5% L-arabinose (w/v), 5 µg/ml tetracycline, and the BAC specific antibiotic. We incubated uREXER cultures grown at 37 °C at 220 rpm until the $OD_{600}$ reached 0.3, at which point cultures were spun down and resuspended into 500 mL of fresh 2xYT media supplemented with 5 µg/ml tetracycline and the BAC specific antibiotic. We then grew cells until the $OD_{600}$ reached 0.5, harvested the cells by centrifugation, and resuspended them in 1 mL of $H_2O$ prior to plating them in serial dilutions on selection plates (245 x 245 x 25 mm, Thermo Scientific). For uREXER selection plates, we supplemented 2xYT agar with 5 µg/ml tetracycline, a negative selection agent against uncut BAC (e.g. 1000 µg/mL streptomycin for odd BACs, or 7.5% (w/v) sucrose for even BACs), a negative selection agent against the upstream genomic landing site (e.g. 1000 µg/mL streptomycin for even genomic landing sites, or 7.5% (w/v) sucrose for odd genomic landing sites), and an antibiotic for the positive downstream marker from the BAC (e.g. 50 µg/mL kanamycin for even BACs, or 20 µg/mL chloramphenicol for odd BACs). We then incubated plates at 37 °C overnight, picked multiple colonies, resuspended them in 50 µL of Milli-Q filtered $H_2O$, and restreaked them onto selective plates. We phenotyped them for uREXER marker swap by arraying 3.5 µL of resuspended colonies on several 2xYT agar plates supplemented with either 50 µg/ml kanamycin, 20 µg/ml chloramphenicol, 1000 µg/ml streptomycin, 7.5% sucrose or 2.5 mM 4-chloro-phenylalanine either on the same day as restreaking, or phenotyped from restreaked colonies. Additionally, the restreaked colonies were again resuspended in 50 µL of Milli-Q filtered $H_2O$ and we genotyped for uREXER marker swap by colony PCR on resuspended colonies using both a primer pair flanking the genomic locus of the upstream landing site and the newly introduced selection cassette from the BAC. Recombinants with successful marker swap yielded a ~500 bp amplicon from the upstream genomic locus (i.e. loss of landing site) and a ~2-3 kb amplicon (i.e. gain of landing site) from the downstream genomic locus. For clones which passed phenotyping and genotyping, we then prepared genomic DNA using the QuickExtract DNA Extraction solution (LGC Biosearch Technologies), according to manufacturer's recommendations, which served as input material for subsequent NGS analyses. All uREXERs were performed as described above, except that the 100k31 full uREXER and the 100k24 NCS library uREXER were performed in a *recA*$^{+}$ MDS42$^{rpsL\text{(K43R)}}$

genomic background. For experiments which required additional negative selection pressure on the genome, we performed REXER4, where four Cas9-based cuts were generated during a synthetic DNA integration experiment – 2 cuts on the BAC to liberate the synthetic DNA as a linear dsDNA substrate, and 2 cuts on the genome flanking the targeted region for integration (**Supplementary Fig. 29**). Additional gRNAs for REXER4 were provided in trans, as described previously[49].

***Synthesis of recoded walk strains by sequential uREXER/GENESIS***

We alternated genomic selection markers for sequential uREXER experiments (GENESIS [49]), enabling the successive integration of 100-kb recoded fragments in a clockwise manner across the genome to thus generate increasingly recoded strains (**Supplementary Fig. 1**). For even BAC uREXERs, we selected against the maintenance of a *sacB-cat* genomic landing site and selected for the integration of a BAC-encoded *rpsL-kanR* cassette. For odd BAC uREXERs, we selected against maintenance of a *rpsL-kanR* genomic landing site and selected for the integration of a BAC-encoded *sacB-cat* cassette.

Following the NGS confirmation of a fully recoded uREXER clone, we inoculated a scrape from the corresponding glycerol stock into 5 mL of 2xYT supplemented with 2% glucose (w/v), 5 µg/mL of tetracycline, and either 50 µg/mL of kanamycin (for even fragments) or 20 µg/mL of chloramphenicol (for odd fragments). After overnight incubation at 37 °C with shaking at 220 rpm, we diluted cultures into larger volumes of 2xYT with the same antibiotic/glucose combination and prepared electrocompetent cells prior to transformation of the next BAC. As described above in the section '***uREXER to integrate 100-kb recoded fragments***', we then proceeded to perform uREXER to integrate the next 100-kb fragment to increasingly recode the walk strain. Typically, GENESIS proceeded for four to five steps (400 to 500 kb) before characterizing recoded strain fitness.

#### *Generation of NCS libraries on a BAC to fix non-recoded regions*

To recode regions that were non-recodable with the vS33A7 recoding scheme, we constructed libraries of small BACs, termed NCS libraries (N-terminal coding sequences), where the 5' ends of coding sequences of selected genes, such as essential or semi-essential genes, were varied. Libraries spanned the first 24-30 bp and preserved the amino acid sequence but varied the synonymous codons, either of recoding events only, or of all codons in that window, and reinstated canonical ATG start codons as needed. The libraries were assembled using degenerate primers (Merck) with which the selected genes were amplified. The entire synthetic insert sequence of the NCS-BAC was fused using overlap extension PCR. The BACs contained the same BAC/YAC backbone as the BACs constructed for uREXER and was amplified from a uREXER BAC with the correct marker setup for the landing site chosen. We assembled the NCS-BACs by standard Gibson assembly protocols, and then electroporated the assembled libraries into *E. coli* MDS42$^{rpsL\text{(K43R)}}$ or MDS42$^{rpsL\text{(K43R)}/\Delta recA}$ cells equipped with the corresponding landing site. We integrated the synthetic sequences into the genome via uREXER and sequenced for fully recoded clones either by Sanger sequencing or NGS.

#### *pTarget protocol for fixing and for walking in 27B*

To recode the 21-kb region within the 64 kb 27B fragment on the genome (**Supplementary Figs. 13, 15**), we first divided this region into 4 pieces, each piece corresponding to the transcriptional unit from one of the 4 main promoters (**Supplementary Fig. 15**). We generated PCR products for each piece using four synthetic DNA templates (Twist Biosciences); these templates are the 21-kb region recoded with four 7-codon compression schemes (vS33A7, vS34A7, vS35A7, or vS36A7). These PCR products were generated with overlap extension PCR for all essential and semi-essential genes (all genes except *rpmJ* and *pioO*) to introduce NCS libraries (and internal promoter libraries for fragment 4) by using primers designed to include synonymisation of the first 8-10 codons and to restore the canonical ATG start codon as needed. Overall, the PCR products varied the recoding across the four schemes (vS33A7, vS34A7, vS35A7, vS36A7) and further varied the NCS and internal promoter sequences. We also split libraries 2 and 3, which contain a large number of genes, into smaller libraries (libraries 5-8) to more effectively cover NCS diversity within these genes (**Supplementary Fig. 15b**). The PCR products were used to replace the corresponding genomic sequence by pTarget-based recombination (CRISPR/Cas9 targeting cleavage of each wild-type genomic sequence and lambda red recombination to integrate the PCR products into the genome[55]).

For pTarget-based recombination, we first transformed a Cas9 plasmid into MDS42$^{rpsL\text{(K43R)}/\Delta recA}$ cells. We designed the pTarget plasmids by duplicating the sgRNA cassettes and replacing the two N20 sequences into new 20-bp sequences that target both ends of each fragment. To recode each fragment, we electroporated the PCR product (5,000-1,000 ng) and the corresponding pTarget plasmid (300-800 ng) simultaneously for targeted genome cutting and homologous recombination. After outgrowth for 3 h, we plated cells under the selection of 50 μg/mL of kanamycin (Cas9 plasmid) and 75 μg/mL of spectinomycin (pTarget). After incubation for 18 hours at 37 °C, we then picked colonies for Sanger sequencing and NGS to confirm the recoding.

After confirming recoding solutions for all 4 fragments individually, we combined the solutions by introducing the sequences into the genome sequentially via pTarget-based recombination using the same protocol (**Supplementary Fig. 15b**). This led to genomic recoding of the 21-

kb region, with a 73 bp wild-type sequence at the C-terminus of *rplP*, which we subsequently replaced by lambda red recombination with a synthetic sequence that implemented the original vS33A7 genome design.

***Individual recoding landscapes and compiled recoding landscapes***

We used an in-house Python script to generate recoding frequency landscapes. Briefly, each recoding event was annotated in the GenBank file using the format [“WT codon” to “Recoded codon”] (e.g. “TCA to AGT”). Next, NGS reads were pre-aligned and sorted by positional index using Bowtie 2[56]. Per sequenced clone, the sorted BAM file was then provided to the script, which scanned through all annotated recoding events in the GenBank file. For each event, the recoding frequency was calculated as the ratio of reads containing the recoded codon to the total number of reads at that position. The script produced individual CSV files for each clone, as well as a final summary CSV file compiling results from all sequenced clones. Resulting landscapes were then visualized in Prism.

***Linkage mapping to determine growth-limiting recoded regions***

In linkage mapping experiments for a given recoded strain, we conjugated genomic regions from a WT donor strain into a recoded recipient strain. The donor strain was prepared by integrating a *gentR-oriT* cassette facing towards the corresponding non-recoded section in the MDS42 genome through lambda red recombineering (described above). The $RK24^{Lux}$ F-plasmid was introduced into the donor strain by electroporation or conjugation, and colonies were obtained on LB agar + 50 µg/mL apramycin and confirmed to contain the $RK24^{Lux}$ F-plasmid by luminescence (BioRad GelDoc, Chemiluminescence, 10 sec). The recipient strain harboured the recoded region of interest flanked by a double-selection cassette as well as the pKW20_CDFSpec_pAra_RecA_Cas9_tracrRNA plasmid. Donor and recipient cells were grown overnight at 37 °C in 2–15 mL of 2xYT selecting for all relevant genomic markers and the F-plasmid, and + 0.5% arabinose for inducing RecA expression in the *ΔrecA* strain. Cells were then harvested by centrifugation (5 min, 5000g) and washed three times with 1 mL of PBS. Each conjugation reaction contained a total of ~$10^9$ cells (estimated by $OD_{600}$) with a 1:1 ratio of donor and recipient cells; multiple replicates were set up in parallel. To initiate the conjugation the mixed cells were pelleted by centrifugation (5 min, 5000g), resuspended in 5–10 µL PBS and spotted on cut cellulose acetate filter membranes (Sartorius, 0.45 µm, Type 11106) placed on TYE/2xYT/LB agar plates (+ 0.5% arabinose for *ΔrecA* strains). The conjugation plates were incubated at 37 °C for ~1.5 h and cells were then recovered by vortexing the membranes in a 2 mL Eppendorf tube with 0.5 mL of SOB. Resuspended cells were then placed in an incubator shaker for 2 h (37 °C, 1,000 rpm). The cells were transferred into 10 mL LB or 2xYT supplemented with 50 µg/mL kanamycin ('even' ending sections) or 20 µg/mL chloramphenicol ('odd' ending sections) and 50 µg/mL spectinomycin, selecting for the recipient cells. The 10 mL cultures were passaged for several days by diluting them 1:100 into fresh media. All passaging intermediates were collected, and cells plated as single colonies ($10^{-6}$-$10^{-7}$ dilution) on TYE/2xYT/LB agar supplemented with 50 µg/mL kanamycin ('even' ending sections) or 20 µg/mL chloramphenicol ('odd' ending sections). From these plates, phenotypically different colonies were picked from TYE plates from all timepoints and replicates into 50 µL of PBS in 1.2 mL 96-well plates (AB-1127). We then added 450 µL of LB containing the appropriate antibiotic to each well and incubated the plates for 24 h (37 °C, 800 rpm). We subjected all picked colonies to: i) growth curves to measure growth parameters, calculated as described in '***Growth rate measurement and analysis***'; ii) NGS analysis, as described in '***Preparation of whole-genome and BAC libraries for next-generation sequencing***'; and iii) recoding landscapes generation, as described in '***Individual recoding landscapes and compiled recoding landscapes***'.

We generated fitness-recoding linkage maps with the recoding and growth data listed above. To identify genomic regions associated with low fitness, we adopted the following strategy: a recoding event that is associated with poor fitness should exhibit a higher doubling time – the strain grows slower when it has not managed to cross out these recoding events across a distribution of different recoding landscapes. We first compiled together growth data and recoding landscapes, collected as described above, before filtering the following: i) samples which gained fitness through mutations outside the recoded region of interest – assumed to have low doubling times/high max $OD_{600}$ but maintain a fully recoded landscape; ii) samples which exhibit 'mixed traces' – assumed to derive from alignments to both a recoded and non-recoded copy of duplicated genome sections; for the purposes of fitness these should be considered non-recoded; and 3) samples with artefactual negative doubling times. Finally, for every recoding event, the minimum, maximum, mean and standard error of doubling times and max $OD_{600}$ were identified, excluding doubling times >1,000 min (assumed non-growing). Linkage maps are plotted as minimum doubling times (black) and mean + stderr of doubling times (red) against corresponding recoding events as genomic coordinates, or maximum Max $OD_{600}$ (black) and mean + standard error of Max $OD_{600}$.

***RNA-Seq mapping in growth limiting regions***

We transformed BACs containing the 100-kb fragment of interest into MDS42$^{\mathit{rpsL}\text{(K43R)}/\Delta \mathit{recA}}$ cells and confirmed their integrity by NGS sequencing as described in ***'Whole-genome and BAC library preparation for Illumina NGS'***. Cells containing the NGS-confirmed BAC were grown overnight in liquid culture in biological triplicates (2xYT + relevant BAC antibiotic); RNA was isolated from diluted overnight culture grown to $OD_{600}$ = 0.2–0.3 (RNeasy Bacteria Mini Kit, Qiagen), and kept on ice during downstream processing. RNA integrity (RIN) scores were assessed on a Tapestation (Agilent) and ensured to be above 7. rRNA was depleted (NEBNext rRNA Depletion Kit (Bacteria), E7860L) to enrich mRNA and libraries were prepared (NEBNext Ultra II Directional RNA Library Prep Kit for Illumina, E7760S) for sequencing on an Illumina NextSeq2000 (P2 XLEAP-SBS (100 cycles), 60 bp PE reads).

Reference files for MDS42$^{rpsL\text{(K43R)}/\Delta recA}$ and each respective BAC were converted from genbank to gff and fasta files using *emboss*[57]. For each BAC, the MDS42$^{rpsL\text{(K43R)}/\Delta recA}$ master gff/fa reference was concatenated with the respective BAC gff/fa. Illumina sequencing reads (FASTQ) were aligned to the 'MDS42 ΔrecA + BAC' reference (*Bowtie 2*,[56]) permitting multiple alignments, and bam files subsequently quantified using *HTSeq*[58]. Reads that aligned to more than one feature (i.e. reads which align to regions of the BAC with no recoding events) are counted for both the genome and the BAC and thus will not be considered differentially expressed.

Read count files were processed using R package *DESEQ2*[59] and transcripts corresponding to regions of the genome outside the 100 kb fragment of interest are excluded. Gene essentiality was annotated according to the Goodall database[60] and Log2FoldChange values plotted against adjusted p-values for each 100kbp fragment using R. Genes outside the range –1<Log2FoldChange <1 were classified as significantly differentially expressed.

***Assembly of a synthetic genome from recoded sections by conjugation***

Recoded sections of semi-synthetic strains were hierarchically combined to yield increasingly recoded strains via conjugative transfer of genomic DNA from a donor and subsequent *recBCD*-mediated recombination in the recipient[61]. Donor strains were prepared by integrating an RK2 *oriT* cassette (*gentR-oriT*, *rpsL-kanR-oriT, sacB-nrsR-oriT*) upstream, and a double-selection cassette downstream, of the synthetic region to be transferred (**Supplementary Fig. 23**). Additionally, the donor strains harboured an RK2 F-plasmid lacking an *oriT* which renders the F-plasmid incapable of self-transfer[51]. The F-plasmid was introduced into the donor strain by electroporation or conjugation. Recipient strains were prepared by integrating a compatible double-selection marker cassette, such as *pheS*- hygR*, downstream of a recoded stretch with homology to the region to be transferred from the donor strain. Regions of homology between donor and recipient were at least 3 kb in length and in some cases this homology was integrated simultaneously with the double-selection cassette integration. Longer homology regions generally led to increased assembly efficiency. Additionally, recipient strains harboured a plasmid encoding for recA under an arabinose-inducible promoter and a positive marker to be selected for after the conjugation, such as *specR*. The expression of recA from the plasmid was induced both in overnight cultures of recipient and in the agar plates on which the conjugation was run when genomic *recA* was lacking. For conjugation, overnight cultures of recipient and donor strains, supplemented with the necessary antibiotics, were washed thrice with PBS and mixed at donor:recipient ratios ranging from 4:1 to 1:1, generally at a total scale of 5 mL of $OD_{600}$ = 2. These mixtures were then centrifuged, and the cell pellet was suspended in a minimal volume of residual PBS and spotted on cellulose acetate filter membranes (Sartorius) placed on a 2xYT, LB, or TYE agar plate. Conjugations were incubated at 37°C for 10 to 120 mins after which cells were recovered from the membranes by vortexing the membrane in SOB. After shaking incubation at 37°C in SOB, for between 2 and 24 h, the cells were plated on 2xYT or LB agar plates supplemented as needed to select for the marker plasmid in the recipient, the positive marker downstream of the synthetic region transferred from the donor, and against the negative marker downstream of the homology region in the recipient. Colonies that grew on these plates were then screened by phenotyping and confirmed by NGS before proceeding to the next assembly step.

***Whole-genome and BAC library preparation for Illumina NGS***

We isolated gDNA or BACs as described in '***uREXER to integrate 100-kb recoded fragments***' and diluted them at least 1:5 in Milli-Q filtered $H_2O$. BACs assembled in yeast were purified as described in '***BAC assembly and delivery***' We performed library preparation with assistance from an automated platform comprising a Biomek FXp (Beckman Coulter) with integrated thermocycler and fluorescence plate reader (Molecular Devices SpectraMAX I3), following protocols either described previously[53] – input DNA is quantified and diluted for the Nextera XT DNA Preparation Kit (Illumina) – or with the NEBNext UltraExpress FS DNA Library Prep Kit (New England Biolabs) without quantification following manufacturer's instructions at ~ 3X reduced volumes: 1) Fragmentation step: input DNA (3 μL), Milli-Q filtered $H_2O$ (2.5 μL), FS enzyme mix (0.3 μL), FS reaction buffer (1.3 μL); 2) Adapter ligation: UltraExpress ligation master mix (8.5 μL), NEBNext Multiplex oligos for Illumina (Unique Dual Index UMI Adapters DNA Sets 1–4) at 1:10 dilution, (1.7 μL); 3) PCR: MTSC master mix (15.5 μL), NEBNext Primer Mix E7397A (1 μL), 5 cycles. Libraries were purified with AMPure XP (Beckman Coulter) according to manufacturer's instructions (23 μL) and eluted with 0.1X TE or Milli-Q filtered $H_2O$ (13 μL). Libraries were subsequently quantified and pooled using Qubit 1X dsDNA HS assay kit (Thermo Fisher Scientific, Q33230) reagents adapted for a fluorescence plate reader.

Pooled libraries were paired-end sequenced on an Illumina NextSeq2000 according to manufacturer's instructions with either NextSeq2000 P1 (100/300 cycles), P2 (100/200 cycles) or P3 (100 cycles) reagent kits, utilising either XLEAP-SBS or v3 SBS chemistry.

***Illumina NGS data analysis***

Reads were demultiplexed onboard (DRAGEN bcl2fastq 3.8.4 or 4.2.7) or from unindexed fastq.gz files with *demuxFQ* (CRUK CI genomics core). Reads were aligned with *Bowtie 2* (v2.5.1) and sorted and indexed with samtools (v1.17) for viewing in IGV (v2.13.2) or landscape generation.

***Growth rate measurement and analysis***

Bacterial clones were grown overnight at 37 °C in 2xYT with the relevant antibiotic. Overnight cultures were diluted 1:100 and monitored for growth in a 200 µL volume in a 96-well plate. Measurements of $OD_{600}$ were taken every 5 min for 18-24 h on a Tecan Microplate reader.

To determine doubling times, we used the AMIGA fitness analysis software[62] to fit an exponential curve to a total of at least 3 independently grown biological replicates. For doubling time and max $OD_{600}$ metrics, the mean and standard deviation from the mean were calculated for all $n \geq 3$ replicates.

**Code Availability**

Scripts for RNA-Seq analysis, recoding landscape generation, and fitness recoding linkage mapping are available at (https://github.com/JWChin-Lab).

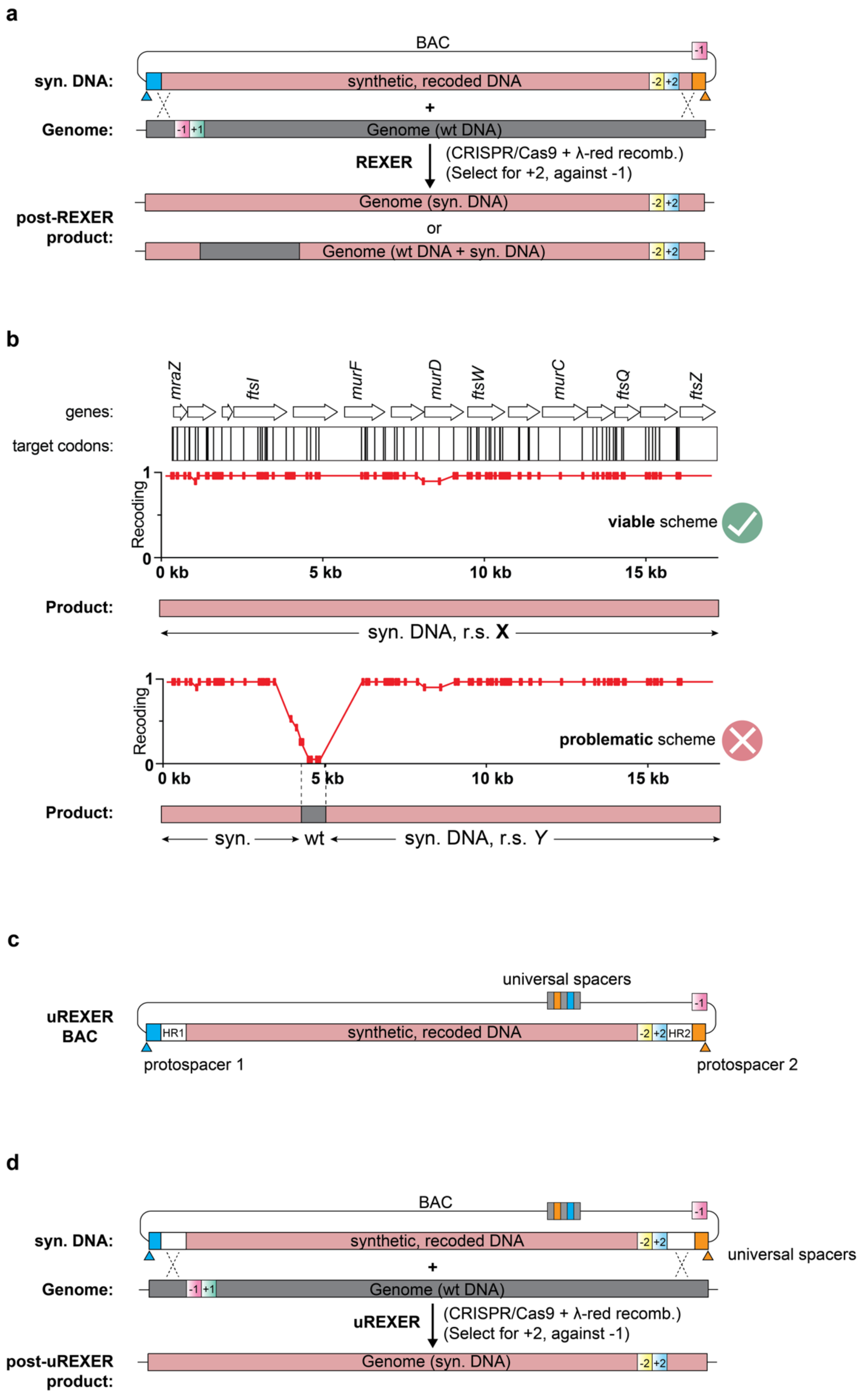


**Supplementary Fig. 1. Using 100-kb fragments of synthetic, recoded DNA to replace the homologous region of the wild type *E. coli* genome and map the allowed and disallowed synthetic sequence via REXER and uREXER.**

**a.** Replicon excision for enhanced genome engineering through programmed recombination (REXER) is an approach which can replace more than 100 kb of the *E. coli* genome with a synthetic DNA sequence in a single step. REXER utilizes i) CRISPR-Cas9 to linearize synthetic DNA provided from an episome (BAC), and ii) lambda-red mediated recombination to replace genomic DNA with the linear synthetic DNA [49]. The light blue and the light orange regions in the BAC denote protospacers which are targeted by CRISPR spacers (light blue and light orange triangles). Induction of Cas9 liberates synthetic DNA (pink) flanked by homology arms to their targeted region of the *E. coli* genome. Lambda-red facilitates recombination between the homologous sequences (hashed exes), with the double-selection cassette (-1/+1) on the wild type genome ensuring removal of the wild type genomic DNA and the (-2/+2) cassette on the BAC ensuring integration of the corresponding synthetic DNA. The example shown here uses -1 as *rpsL*, +1 as *KanR*, -2 as *sacB*, and +2 as *CmR*.
**b.** Compiled recoding landscapes plot the extent of recoding at each targeted codon across the population of post-REXER clones. In this example, REXER is used to replace wt genomic DNA with recoded DNA over a 20-kb genomic region rich in essential genes and target codons. Post-REXER clones are subject to next-generation sequencing (NGS). The compiled recoding landscape plots the frequency at which each target codon is recoded across the population of clones. For a viable recoding scheme the recoding frequency in the compiled recoding landscape never goes to zero, and individual fully recoded clones are identified. For non-viable schemes the compiled recoding landscape identifies regions where the recoding frequency goes to zero and no fully recoded clones are identified; for these schemes the compiled recoding landscape identifies the of wt DNA that are recalcitrant to recoding using this scheme.
**c.** Universal REXER (uREXER) utilizes universal CRISPR/Cas9 spacers encoded from the BAC to facilitate the integration of 100-kb of synthetic DNA in a single step. The light blue and the light orange rectangular regions in the BAC denote the universal protospacers targeted by the corresponding universal spacers cassette (light blue and light orange squares).
**d.** Universal spacers enable linearization of the BAC following induction of Cas9 activity in the cell. The synthetic DNA is then integrated into the genome via the homology regions located on the 5' and 3' ends of the BAC (as described above in panel **a**).

**a**

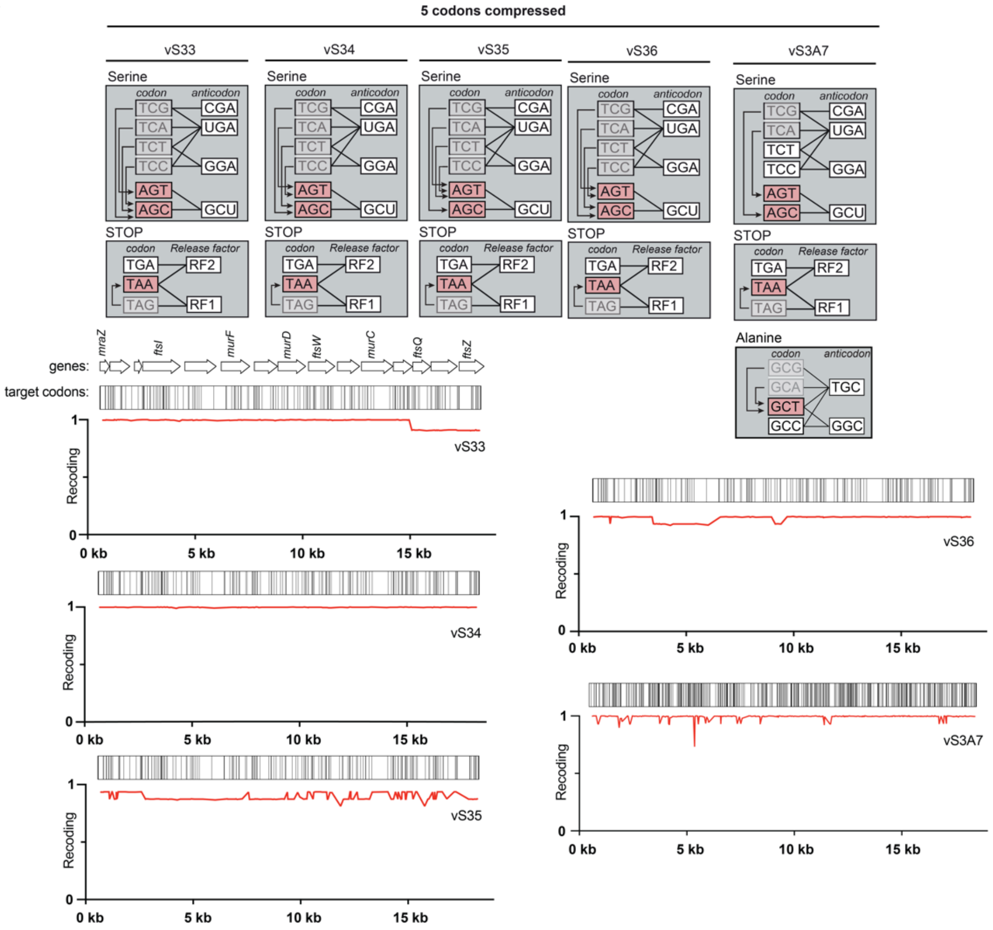
5 codons compressed
vS33
vS34
vS35
vS36
vS3A7
Serine
codon
anticodon
TCG
TCA
TCT
TCC
AGT
AGC
CGA
UGA
GGA
GCU
STOP
Release factor
TGA
TAA
TAG
RF2
RF1
Alanine
GCG
GCA
GCT
GCC
TGC
GGC
genes:
target codons:
mraZ
ftsI
murF
murD
ftsW
murC
ftsQ
ftsZ
Recoding
0 kb
5 kb
10 kb
15 kb

**b**

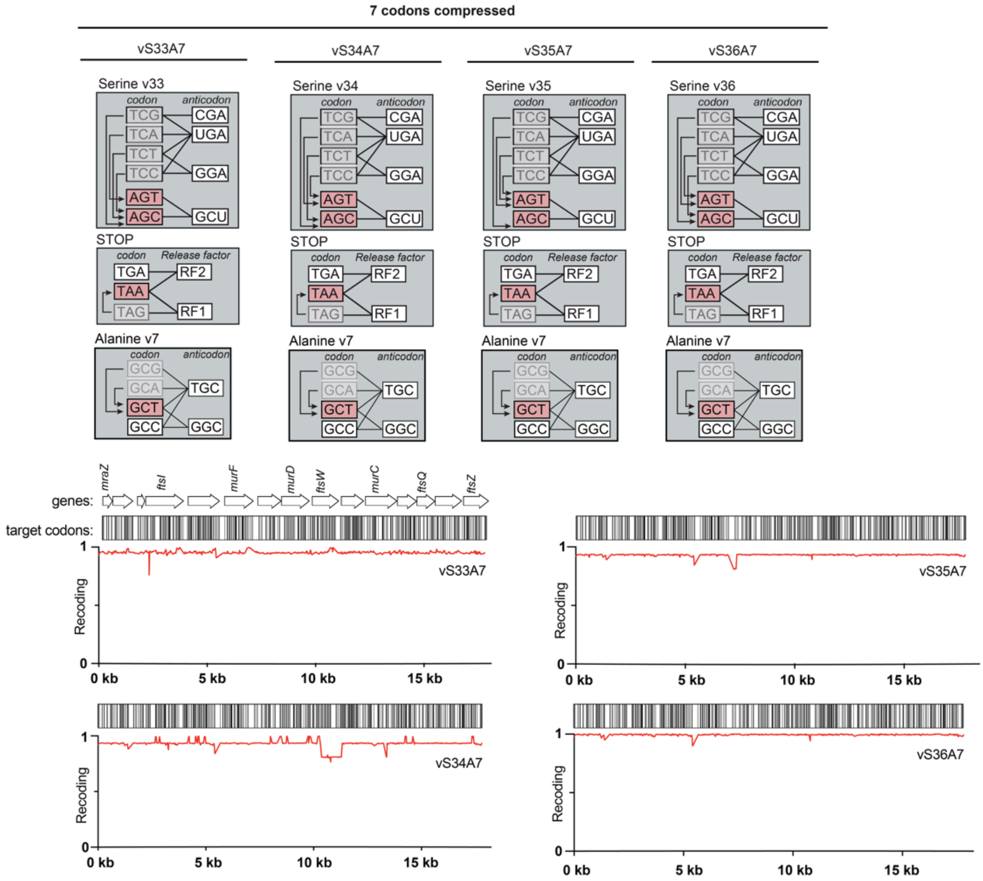
7 codons compressed
vS33A7
vS34A7
vS35A7
vS36A7
Serine v33
Serine v34
Serine v35
Serine v36
Alanine v7
codon
anticodon
STOP
Release factor
genes:
target codons:
Recoding
0 kb
5 kb
10 kb
15 kb

**Supplementary Fig. 2. Compiled recoding landscapes for combining codon compression schemes to provide a 57-codon genome design.**

**a.** Compressing the genetic code by 5 codons via recoding a total of 4 sense codons and 1 stop codon. The schemes recode either: i) 2 serine codons, 2 alanine codons, and 1 stop codon; or ii) 4 serine codons and 1 stop codon. To experimentally evaluate the success of different codon compression schemes, we performed REXER to replace wt with recoded DNA over a test 20-kb genomic region (grey) rich in essential genes and target codons [49]. Post-REXER clones were subject to next-generation sequencing (NGS) and the data used to generate compiled recoding schemes for the 5-codon compression schemes, vS33, vS34, vS35, vS36, and vS3A7. These schemes yielded successful recoding across the 20-kb test region demonstrating that different combinations of sense and stop recoding schemes could be combined. All compiled recoding landscapes were generated from 16 sequenced clones.

**b.** Compressing the genetic code by 7 codons via recoding a total of 6 sense codons and 1 stop codon. We tested four different 7-codon schemes, vS33A7, vS34A7, vS34A7, and vS36A7. These schemes recode different combinations of 4 serine codons, 2 alanine codons, and 1 stop codon. Compiled recoding landscapes show that all four schemes enabled full-recoding across the 20-kb test region. The recoding landscape for vS33A7 is also shown in **Fig. 1** – this scheme was chosen to implement across the entire 4 Mb *E. coli* template genome to provide the initial design for Syn57. All compiled recoding landscapes were generated from 16 sequenced clones.

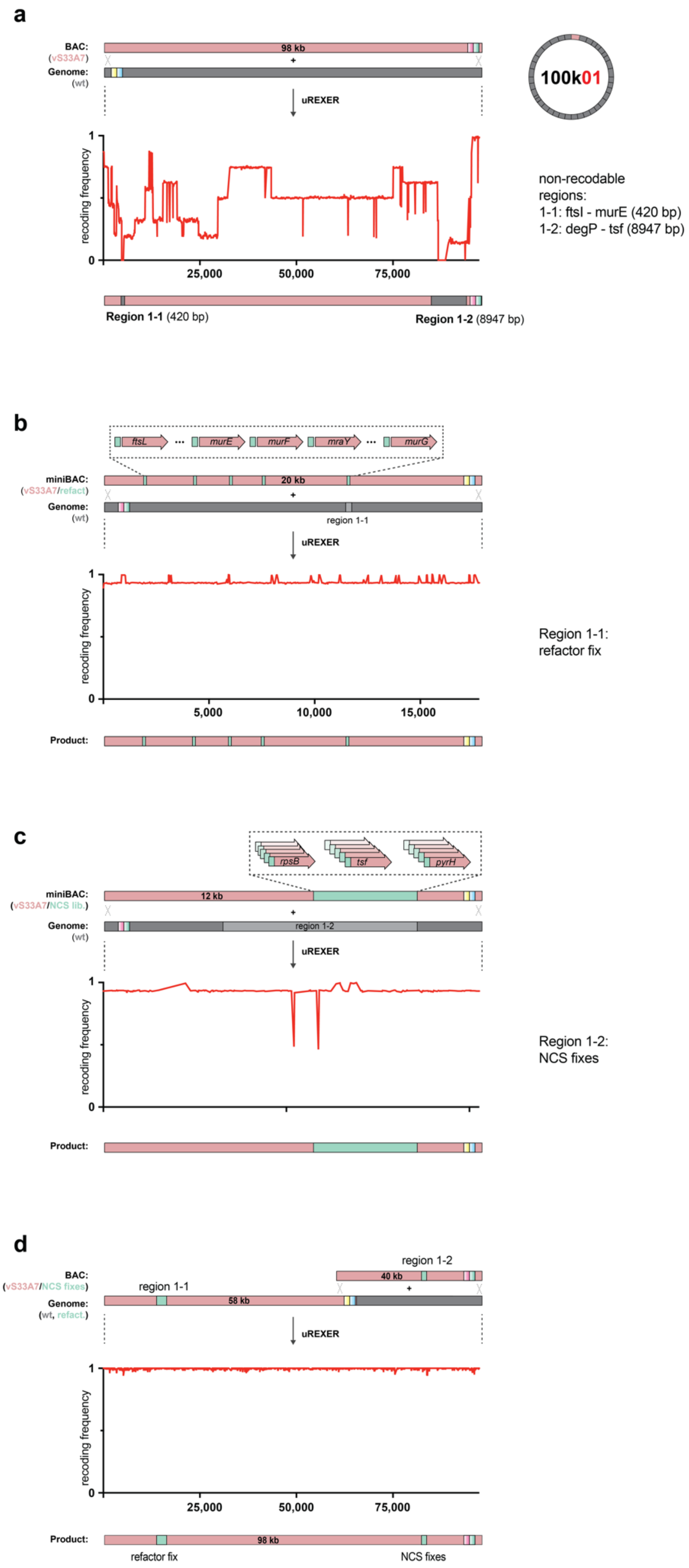
a
BAC:
(vS33A7)
98 kb
Genome:
(wt)
uREXER
100k01
recoding frequency
25,000
50,000
75,000
non-recodable regions:
1-1: ftsI - murE (420 bp)
1-2: degP - tsf (8947 bp)
Region 1-1 (420 bp)
Region 1-2 (8947 bp)
b
ftsL
murE
murF
mraY
murG
miniBAC:
(vS33A7/refact)
20 kb
Genome:
(wt)
region 1-1
uREXER
recoding frequency
5,000
10,000
15,000
Region 1-1:
refactor fix
Product:
c
rpsB
tsf
pyrH
miniBAC:
(vS33A7/NCS lib.)
12 kb
Genome:
(wt)
region 1-2
uREXER
recoding frequency
Region 1-2:
NCS fixes
Product:
d
region 1-2
BAC:
(vS33A7/NCS fixes)
40 kb
region 1-1
Genome:
(wt, refact.)
58 kb
uREXER
recoding frequency
25,000
50,000
75,000
Product:
98 kb
refactor fix
NCS fixes

**Supplementary Fig. 3. Fully recoding 100k01 in the genome.**

**a.** Compiled recoding landscape (from 8 clones) from the 100k01 uREXER. The synthetic, recoded, DNA from the BAC (pink) was used for uREXER into the wild-type (grey) genome. The resulting compiled recoding landscape revealed two regions on the genome which were not recoded in any clone: region 1-1 was 420 bp and region 1-2 was 6.7 kb. The essential genes *rpsB* and *tsf* sit at the 3' end of region 1-2– for the purposes of fixing we expanded region 1-2 beyond the region mapped to include these essential genes; the expanded 1-2 region spanned 8.9 kb.

**b.** A miniBAC containing 20 kb of synthetic recoded DNA covering region 1-1 of the genome, which was not recoded in the initial uREXER in **a**, was used for a new uREXER. The miniBAC used the same recoding scheme as the 98-kb BAC for 100k01. However, the miniBAC harboured larger synthetic inserts for the refactoring events between the *mraW* and *ftsL* genes, the *ftsI* and *murE* genes, the *murE* and *murF* genes, and the *murF* and *mraY* genes. Following REXER with the refactoring fix miniBAC, the resulting compiled recoding landscape (from 16 clones) led to full recoding in the 100k01 fragment region containing region 1-1.

**c.** A miniBAC library containing 12 kb of synthetic recoded DNA covering region 1-2 of the genome that was not recoded in the initial uREXER in **a** and adjacent essential genes was created and used for a new uREXER. This miniBAC contained an NCS library (turquoise) that varied the codons for the N-terminal sequences of *rpsB*, *tsf* and *pyrH.* uREXER with this miniBAC NCS library led to full recoding of the corresponding genomic region – the recoding landscape of the final clone shows full recoding across region 1-2.

**d.** We assembled another miniBAC harbouring the recoded sequence variant identified in **c**. The resulting miniBAC was used for uREXER on top of a genomic template containing the refactor fix in region 1-1 (from panel **b**) and recoded sequence up until region 1-2 - the resulting recoding landscape from a single clone shows full recoding across the entire 98-kb fragment of 100k01.

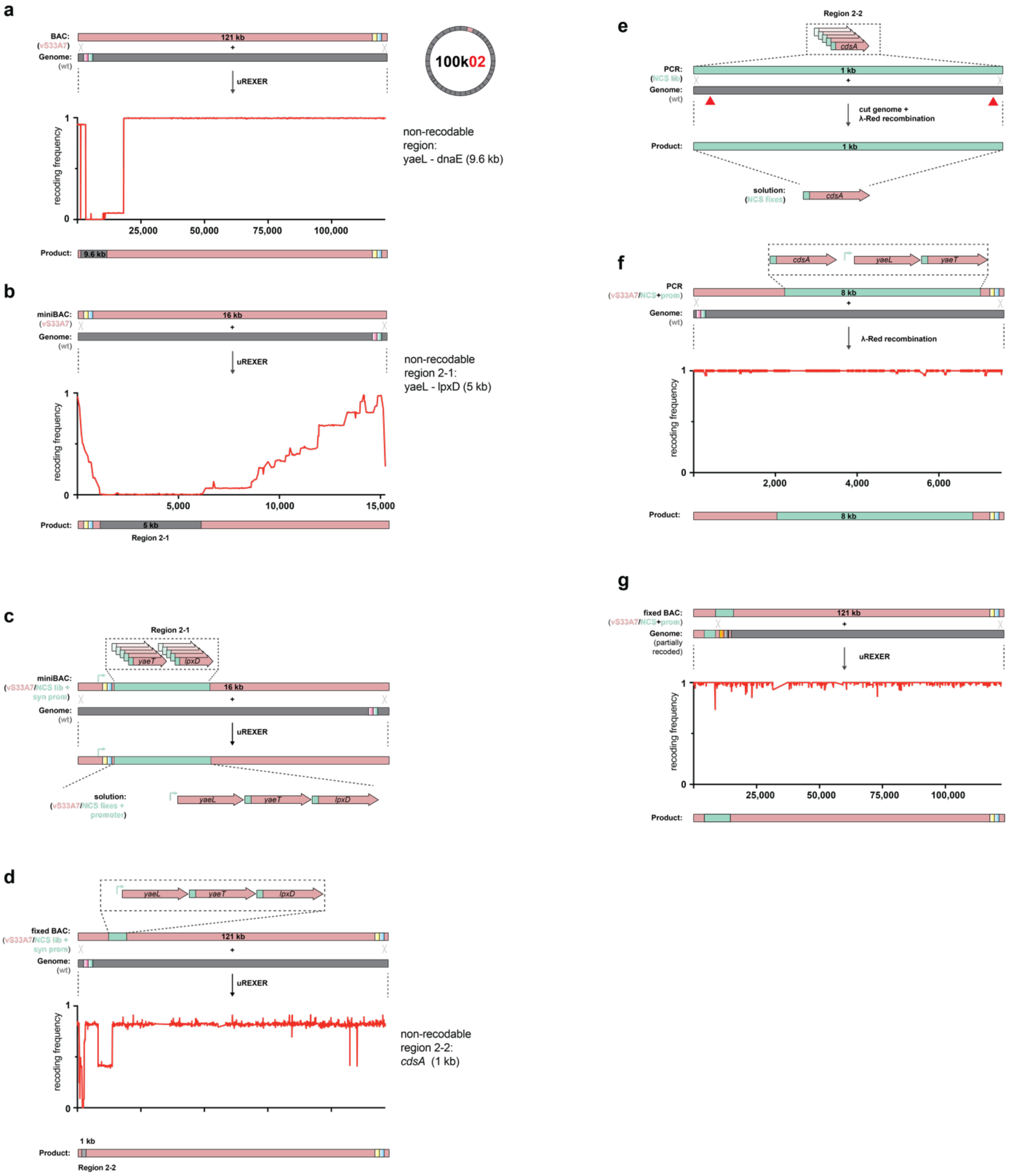


**Supplementary Fig. 4. Fully recoding 100k02 in the genome.**

**a.** Compiled recoding landscape (from 32 clones) from the 100k02 uREXER. The synthetic, recoded, DNA from the BAC (pink) was used for uREXER into the wild-type (grey) genome. The resulting compiled recoding landscape revealed a 9.6 kb region of the genome, from *yaeL* to *dnaE*, which was not recoded.

**b.** A miniBAC containing the synthetic recoded DNA covering the region of the genome that was not recoded in the initial uREXER was used for a new uREXER. The resulting compiled recoding landscape (from 16 clones) refined the region of the genome that was not recoded to 5 kb (grey) spanning from *yaeL* to *lpxD* (region 2-1).

**c.** A miniBAC library containing the synthetic recoded DNA covering the region of the genome that was not recoded in the uREXER in **c** was created and used for a new uREXER. The miniBAC contained an NCS library for the *yaeT* and *lpxD* genes.
**d.** The sequence variant identified in **c** was incorporated into the BAC for 100k02, and the resulting BAC was used for uREXER. The resulting compiled recoding landscape (from 8 clones) is shown. This experiment identified a 1-kb region (region 2-2, grey) within the *cdsA* gene that remains non-recoded in the genome.
**e.** A PCR product containing an NCS library of *cdsA* was used as an input for a lambda red recombination experiment with CRISPR-Cas9 directed genome cleavage (red triangles) enabled the recoding of the *cdsA* gene.
**f.** Solutions identified in previous steps (turquoise for NCS and yellow for promoter) were integrated into one PCR product and used as the input for the lambda red recombination. This led to full recoding of the first 8 kb of 100k02 on the genome, with the recoded genomic region including regions 2-1 and 2-2. The data shows the recoding landscape of the individual clone taken forward for future experiments.
**g.** The genome generated in **f** provided a template for uREXER with the BAC used in **d.** This led to complete recoding of 100k02 on the genome. The data shows the compiled recoding landscape (from 48 clones).

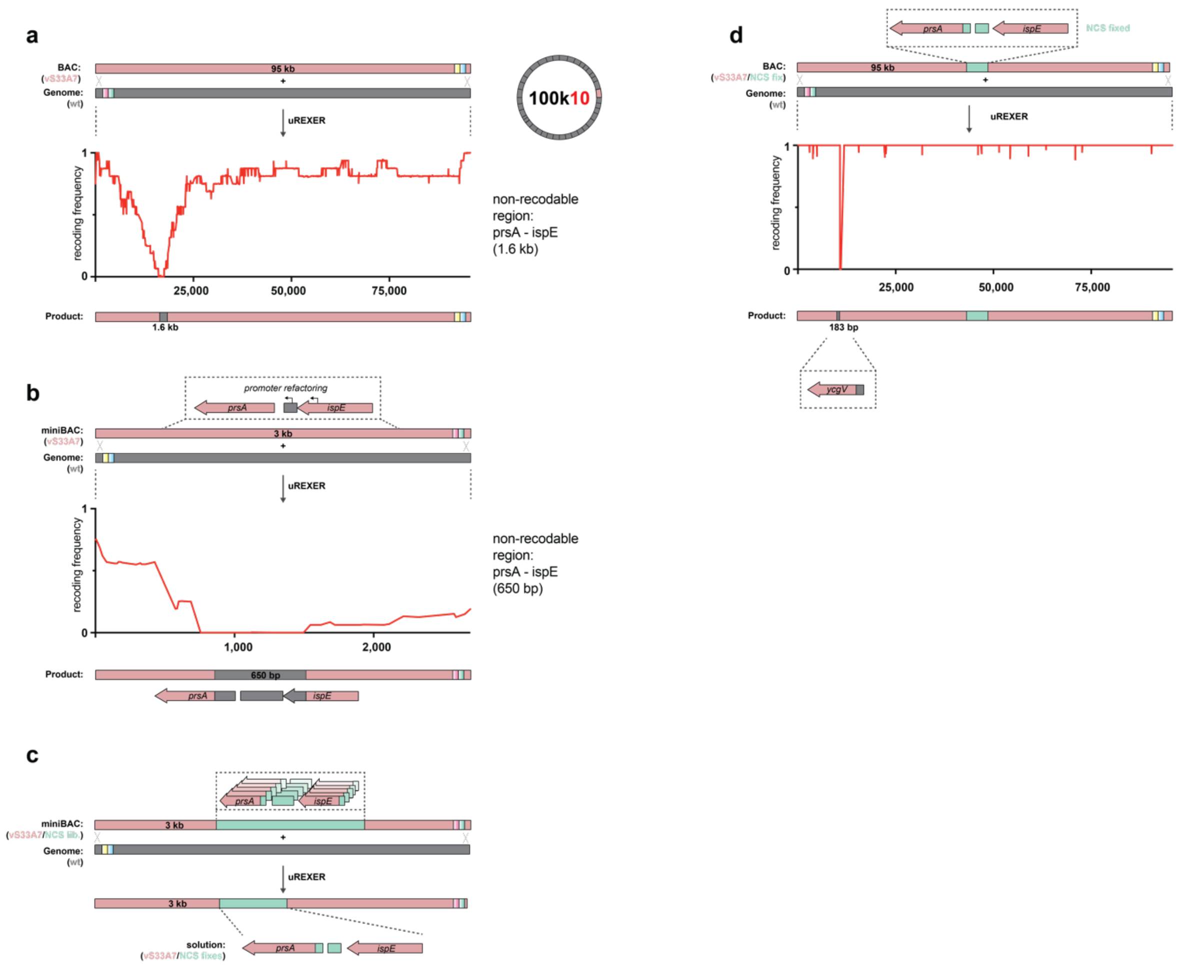


**Supplementary Fig. 5. Fully recoding 100k10 in the genome**.

**a.** Compiled recoding landscape (from 16 clones) from the 100k10 uREXER. The synthetic, recoded, DNA from the BAC (pink) was used for uREXER into the wild-type (grey) genome. The resulting compiled recoding landscape revealed a 1.6 kb region of the genome, from *prsA* to *ispE*, which was not recoded.

**b.** A miniBAC containing the synthetic recoded DNA covering the region of the genome that was not recoded in the initial uREXER was used for a new uREXER; the miniBAC contained additional refactoring of the *prsA* promoter which lies within *ispE* (dotted box). The resulting compiled recoding landscape (from 16 clones) refined the region of the genome that was not recoded to 650 bp (grey).

**c.** A miniBAC library containing the synthetic recoded DNA covering the region of the genome that was not recoded in the uREXER in **b** was created. The miniBAC contained an NCS library (turquoise) that varied the codons for the N-terminal sequence of *prsA* and *ispE* as well as parts of the intergenic region (dotted box).

**d.** The 100k10 BAC used in **a** was modified with the sequence discovered in **c** and the resulting BAC was used for uREXER. The only correctly assembled BAC contained 6 wild-type codons spanning a region of 183 bp. Since this region was previously recoded, we proceeded with the uREXER. The recoding landscape of the correct clone shows full recoding across the entire fragment, with the exception of the 6 wild-type codons that were already present on the BAC. These 6 wild-type codons are within the gene *ycgV*, as shown. This 183 bp wild-type region from was later fixed via lambda-red recombination to yield full recoding across the entire 100k10 fragment.

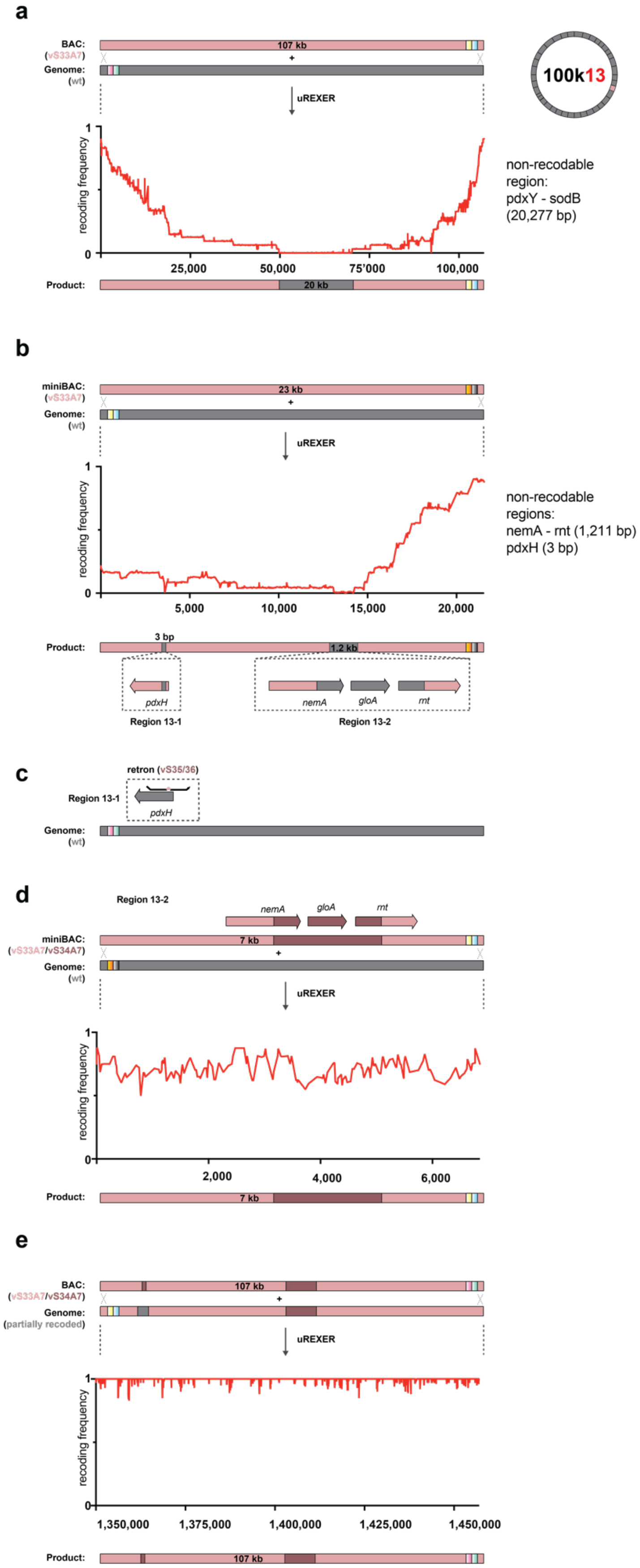
a
BAC:
(vS33A7)
Genome:
(wt)
107 kb
uREXER
100k13
recoding frequency
25,000
50,000
75'000
100,000
Product:
20 kb
non-recodable region:
pdxY - sodB
(20,277 bp)
b
miniBAC:
(vS33A7)
Genome:
(wt)
23 kb
uREXER
recoding frequency
5,000
10,000
15,000
20,000
non-recodable regions:
nemA - rnt (1,211 bp)
pdxH (3 bp)
Product:
3 bp
1.2 kb
pdxH
nemA
gloA
rnt
Region 13-1
Region 13-2
c
retron (vS35/36)
Region 13-1
pdxH
Genome:
(wt)
d
Region 13-2
nemA
gloA
rnt
miniBAC:
(vS33A7/vS34A7)
Genome:
(wt)
7 kb
uREXER
recoding frequency
2,000
4,000
6,000
Product:
7 kb
e
BAC:
(vS33A7/vS34A7)
Genome:
(partially recoded)
107 kb
uREXER
recoding frequency
1,350,000
1,375,000
1,400,000
1,425,000
1,450,000
Product:
107 kb

**Supplementary Fig. 6. Fully recoding 100k13 in the genome.**

**a.** Compiled recoding landscape (from 16 clones) from the 100k13 uREXER. The synthetic, recoded, DNA from the BAC (pink) was used for uREXER into the wild-type (grey) genome. The resulting compiled recoding landscape revealed a 20.3 kb region of the genome, from *pdxY* to *sodB*, which was not recoded. Additionally, we observed a single isolated non-recoded codon, which we did not further investigate.

**b.** A miniBAC containing the synthetic recoded DNA covering the region of the genome that was not recoded in the initial uREXER was used for a new uREXER. The resulting compiled recoding landscape (from 25 clones) refined the region of the genome that was not recoded to 1.2 kb (grey) spanning from *nemA* to *rnt* (dotted box, region 13-2), and an isolated region of 3 bp (Ser codon) in *pdxH* (dotted box, region 13-1).

**c.** To assess an alternative recoding scheme for the Ser codon at the start of *pdxH* that was not recoded in the genome (region 13-1), we edited this codon on the wt genome. This revealed a TCT to AGT substitution to be viable.

**d.** A miniBAC containing synthetic DNA spanning region 13-2 was constructed, the synthetic DNA used the alternative recoding scheme vS34A7 (maroon) for the parts of *nemA*, *gloA*, and *rnt* that were not recoded on the genome using the original genome design. uREXER with this miniBAC led to full recoding of the corresponding genomic region. The compiled recoding landscape (from 16 clones) shows full recoding across the entire fragment.

**e.** The 100k10 BAC used in **a** was modified with the sequences discovered in **c** and **d** and the resulting BAC was used for uREXER on a previously partially recoded clone. The recoding landscape from the clone taken forward shows full recoding across the entire fragment.

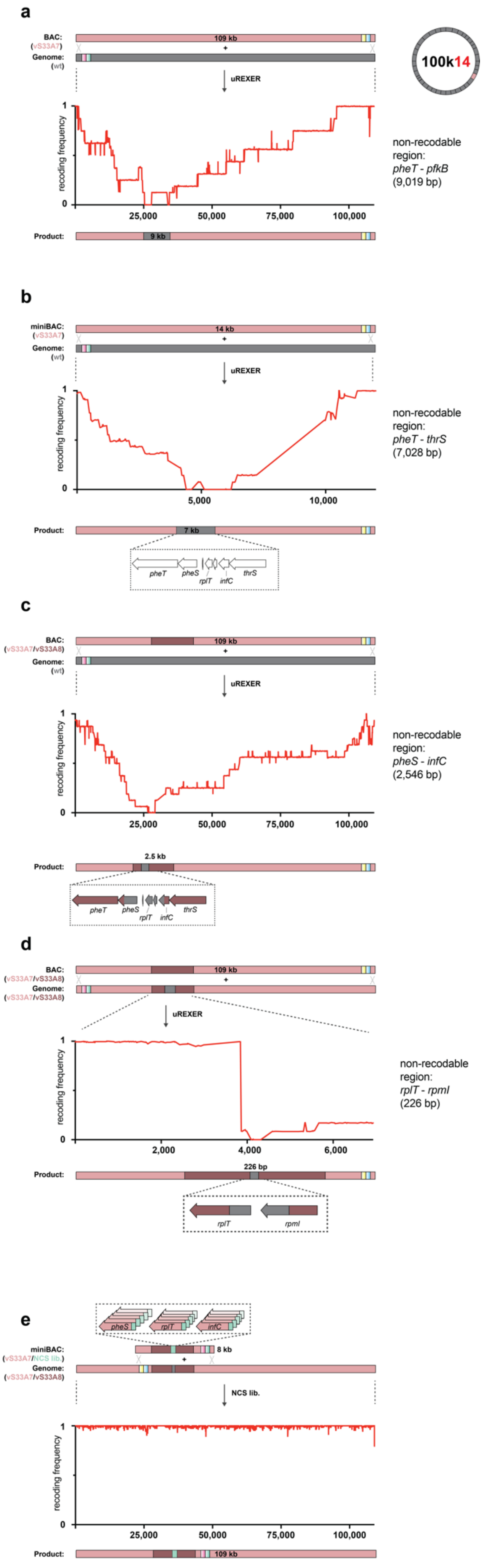
a
BAC:
(vS33A7)
Genome:
(wt)
109 kb
100k14
uREXER
recoding frequency
25,000
50,000
75,000
100,000
Product:
9 kb
non-recodable region: pheT - pfkB (9,019 bp)
b
miniBAC:
(vS33A7)
Genome:
(wt)
14 kb
uREXER
recoding frequency
5,000
10,000
Product:
7 kb
pheT
pheS
rplT
infC
thrS
non-recodable region: pheT - thrS (7,028 bp)
c
BAC:
(vS33A7/vS33A8)
Genome:
(wt)
109 kb
uREXER
recoding frequency
25,000
50,000
75,000
100,000
Product:
2.5 kb
pheT
pheS
rplT
infC
thrS
non-recodable region: pheS - infC (2,546 bp)
d
BAC:
(vS33A7/vS33A8)
Genome:
(vS33A7/vS33A8)
109 kb
uREXER
recoding frequency
2,000
4,000
6,000
Product:
226 bp
rplT
rpmI
non-recodable region: rplT - rpmI (226 bp)
e
pheS
rplT
infC
miniBAC:
(vS33A7/NCS lib.)
Genome:
(vS33A7/vS33A8)
8 kb
NCS lib.
recoding frequency
25,000
50,000
75,000
100,000
Product:
109 kb

**Supplementary Fig. 7. Fully recoding 100k14 in the genome.**

**a.** Compiled recoding landscape (from 16 clones) from the 100k14 uREXER. The synthetic, recoded, DNA from the BAC (pink) was used for uREXER into the wild-type (grey) genome. The resulting compiled recoding landscape revealed a 9 kb region of the genome, from *pheT* to *pfkB*, which was not recoded.

**b.** A miniBAC containing the synthetic recoded DNA covering the region of the genome that was not recoded in the initial uREXER was used for a new uREXER. The resulting compiled recoding landscape (from 16 clones) refined the region of the genome that was not recoded to 7 kb (grey) spanning from *pheT* to *thrS*.

**c.** To assess an alternative recoding scheme for the region between *pheT* to *thrS* that was not recoded on the genome, we assembled a 109-kb BAC harbouring the vS33A7 scheme throughout except with the vS33A8 alternative recoding scheme for genes *pheT* to *thrS*. REXER with this alternative BAC yielded a compiled recoding landscape (from 48 clones) that refined the region of the genome that was not recoded to 2.5 kb (grey).

**d.** Repeating the REXER with the alternative recoding scheme, except this time on to a genomic template from the previous experiment in panel **c**, yielded a compiled recoding landscape (from 12 clones) that refined the region of the genome that was not recoded to 226 bp between the *rplT* and *rpmI* genes (grey).

**e.** A miniBAC containing synthetic DNA spanning the *pheS*, *rplT*, and *infC* genes was constructed, in which the synthetic DNA contained an NCS library of these three genes. uREXER with this miniBAC into the genome of the most recoded clone generated in panel **d** led to full recoding of the corresponding genomic region – the recoding landscape of the final clone shows full recoding across the entire 100k14 fragment.

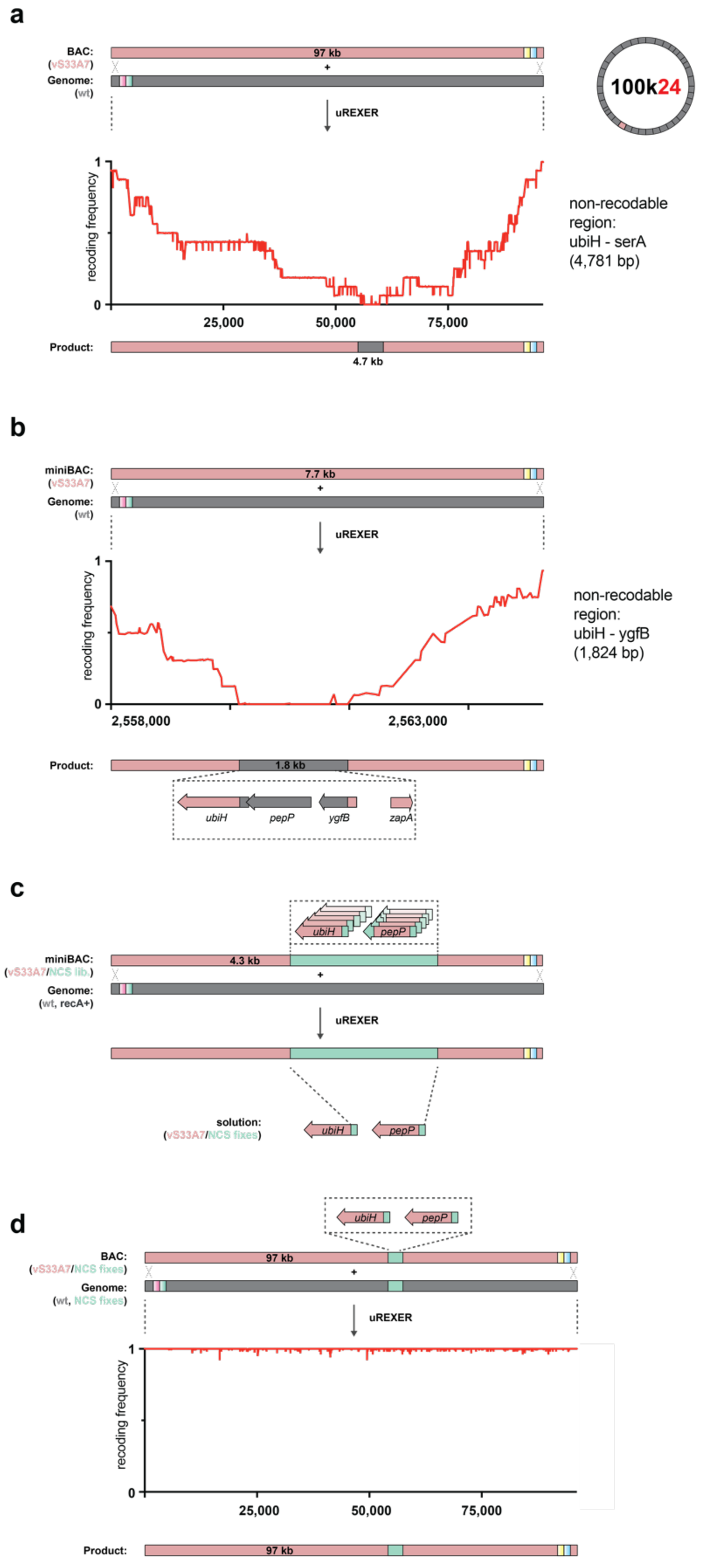


**Supplementary Fig. 8. Fully recoding 100k24 in the genome.**
**a.** Compiled recoding landscape (from 16 clones) from the 100k24 uREXER. The synthetic, recoded, DNA from the BAC (pink) was used for uREXER into the wild-type (grey) genome. The resulting compiled recoding landscape revealed a 4.8 kb region of the genome, from *ubiH* to *serA*, which was not recoded.

**b.** A miniBAC containing the synthetic recoded DNA covering the region of the genome that was not recoded in the initial uREXER was used for a new uREXER. The resulting compiled recoding landscape (from 16 clones) refined the region of the genome that was not recoded to 1.8 kb (grey) spanning *ubiH* to *ygfB* (dotted box).
**c.** A miniBAC library containing the synthetic recoded DNA covering the region of the genome that was not recoded in the uREXER in **b** was created and used for a new uREXER. The miniBAC contained an NCS library (dotted box) that varied the codons for the N-terminal sequence of *ubiH* and *pepP* (turquoise) and a C-terminal region of *pepP* which is hypothesized to contain the RBS for *ubiH*.
**d.** The 100k24 BAC used in **a** was modified with the sequence discovered in **c** and the resulting BAC was used for uREXER on a clone that already contained the solution for *ubiH* and *pepP*. The recoding landscape of the correct clone shows full recoding across the entire fragment.

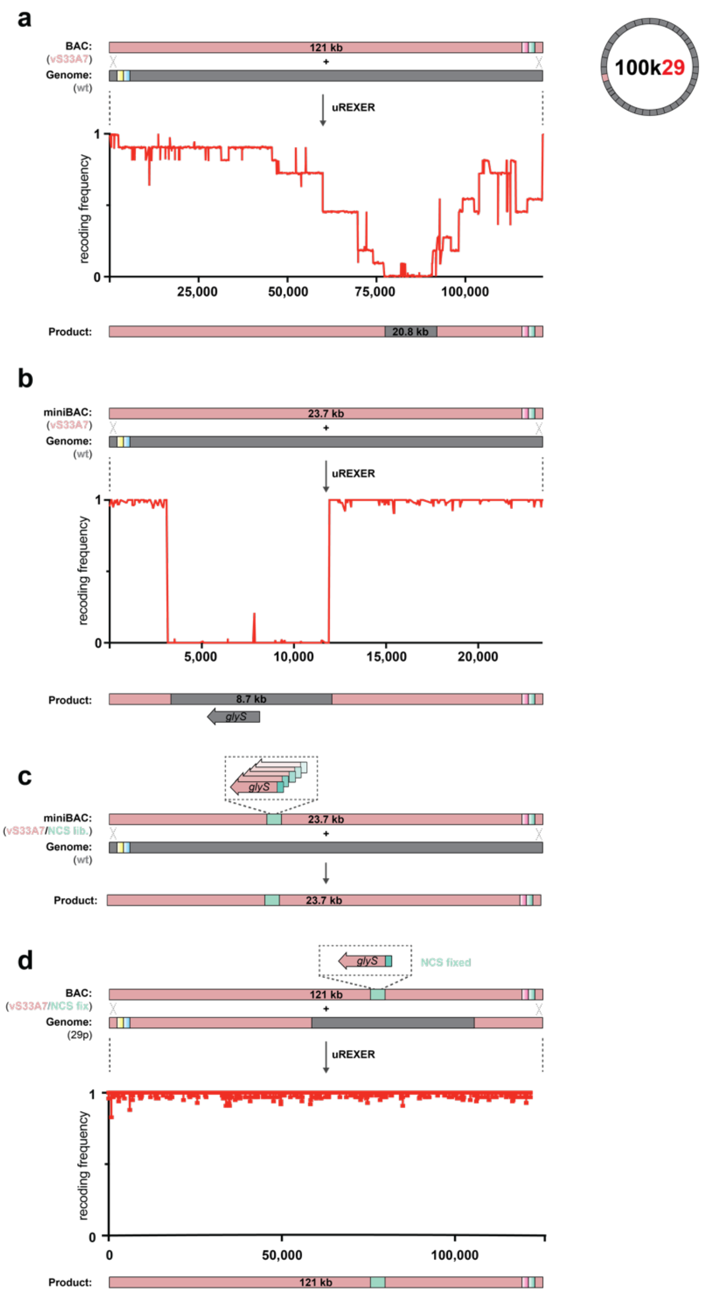


**Supplementary Fig. 9. Fully recoding 100k29 in the genome.**
**a.** Compiled recoding landscape (from 11 clones) of the 100k29 uREXER. The BAC (pink) was integrated into the wild-type (grey) genome. The recoding landscape revealed a non-recoded region of 20.8 kb.
**b.** A miniBAC was constructed spanning the non-recoded region. The recoding landscape from the most recoded clone refined the non-recoded region to 8.7 kb.
**c.** For the refined non-recoded region, a NCS library was constructed (turquoise) in a mini-BAC. The NCS library miniBAC was used for uREXERinto a wt genome; this identified an NCS solution for *glyS* which enabled the integration of the complete synthetic DNA sequence from the mini-Bac into the genome.
**d.** The NCS solution identified in panel **c** was incorporated into the full 100k29 BAC. The modified 100k29 BAC was used for uREXER into a partially recoded genome (29p), obtained

from the experiment described in panel **a**. The recoding landscape from a resulting fully recoded clone is shown.

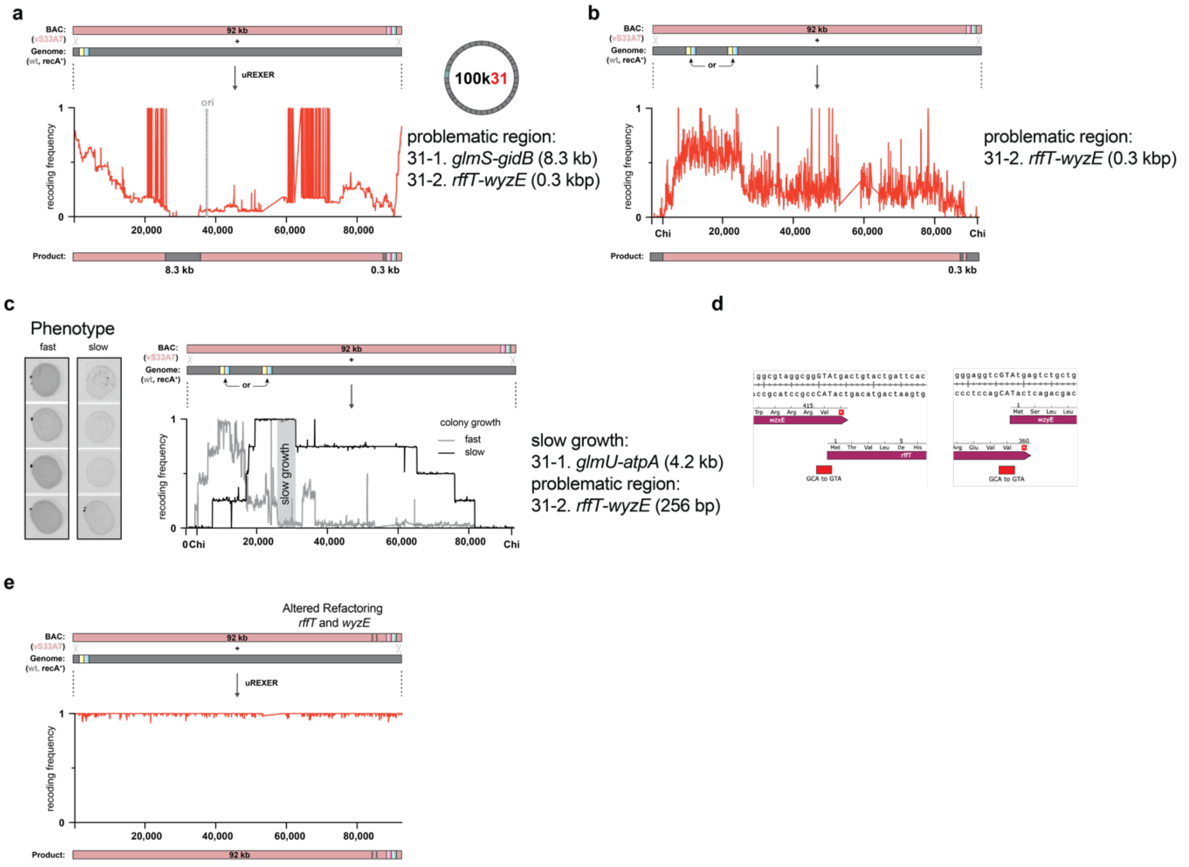


**Supplementary Fig. 10. Fully recoding 100k31 in the genome.**

**a.** Compiled recoding landscape (from 24 clones) from the 100k31 uREXER. The synthetic, recoded, DNA from the BAC (pink) was used for uREXER into the wild-type (grey) genome; these experiments were performed in RecA positive cells. The resulting compiled recoding landscape revealed two regions of the genome which were not recoded: region 31-1 is 35.3 kb, region 31-2 is 0.3 kb.

**b.** Recombinants were selected for via loss of a genomic marker placed at one of the two locations indicated in the genome. The compiled recoding landscape shows the recoding frequency for a pool of 48 colonies (24 for each site). This indicated that the problematic region 31-1 can be fully recoded, while 31-2 remains non-recoded.

**c.** The recombinants obtained in **b** showed either fast or slow growth. By comparing the compiled recoding landscape from four representative clones of each phenotype we mapped the slow growth phenotype to a 4.2 kb genomic section (*glmU* to *atpA*, grey box).

**d.** The problematic region 31-2 is localized to the *wzxE* and *rffT* locus. In the initial genome design, both genes were refactored through introduction of a 20 bp insertion. In both cases this was done due to the overlap of their respective start codon with a recoding event at the last position of their preceding gene. To reinstate the overlap, we mutated the alanine codon (GCA) to valine (GTA), effectively removing the target codon.

**e.** The design change described in **d** was incorporated into the 100k31 BAC used in **a** and the resulting BAC was used for uREXER. We picked slow growing colonies which only become visible after two or more days of incubation. This led to the identification of clones that were recoded across the entire fragment, and the recoding landscape of one such clone is shown.

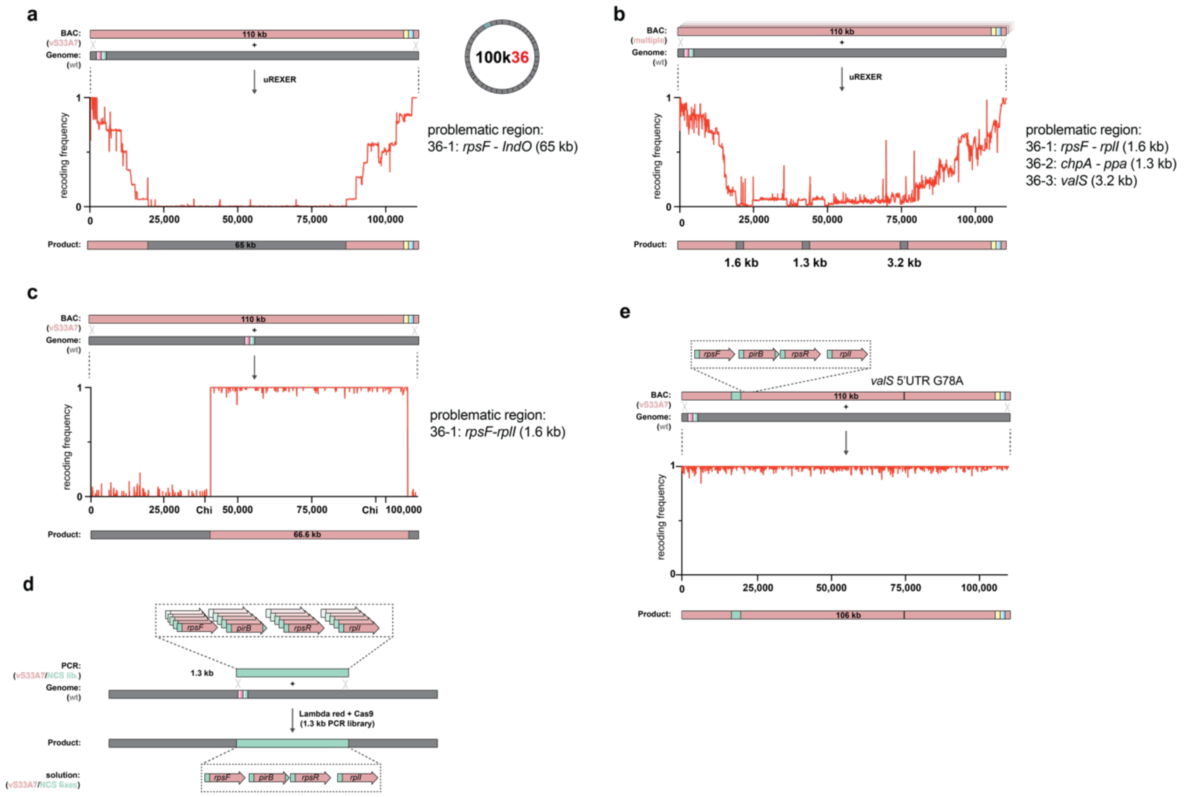


**Supplementary Fig. 11. Fully recoding 100k36 in the genome.**

**a.** Compiled recoding landscape (from 15 clones) from the 100k36 uREXER. The synthetic, recoded, DNA from the BAC (pink) was used for uREXER into the wild-type (grey) genome. The resulting compiled recoding landscape revealed a 65-kb region of the genome which was not recoded.

**b.** uREXER experiments were performed with two variants of the 100k36 BAC. In these BACs all essential (*rpsF*, *pirB*, *rpsR*, *ppa*, *valS*) and semi-essential genes (*chpAB*, *rplI*) within the 65-kb region defined in **a** were recoded using the alternative recoding scheme vS35A7 or vS33A10 while the remaining sequence was recoded using vS33A7. A compiled recoding landscape was generated from the sequences of 16 post REXER clones (eight post REXER clones from experiments with each BAC). The alternative recoding schemes did not enable recoding of the essential genes, but this experiment did further refine the problematic region to the following three regions containing these essential genes: region 36-1, 1.6 kb; region 36-2, 1.3 kb; region 36-3, 3.2 kb.

**c.** The recoding landscape shows the most recoded clone obtained through this method. This clone recoded 66.6 kb of genomic sequence covering problematic regions 36-2 and 36-3.

**d.** For region 36-1, an NCS library was constructed (turquoise) by PCR. The library varied the N-terminal codons of *rpsF*, *pirB*, *rpsR* amd *rplI* as well as a C-terminal recoding event within *pirB*. The NCS library was integrated into a wt genome via pTarget-based recombination (CRISPR/Cas9 targeting cleavage of each wild-type genomic sequence and lambda-red recombination) and led to identification of solutions for each gene.

**e.** The NCS library solution and a spontaneous *valS* 5'UTR mutation were incorporated into a 100k36 BAC. The recoding landscape from a clone that is fully recoded across the entire fragment is shown.

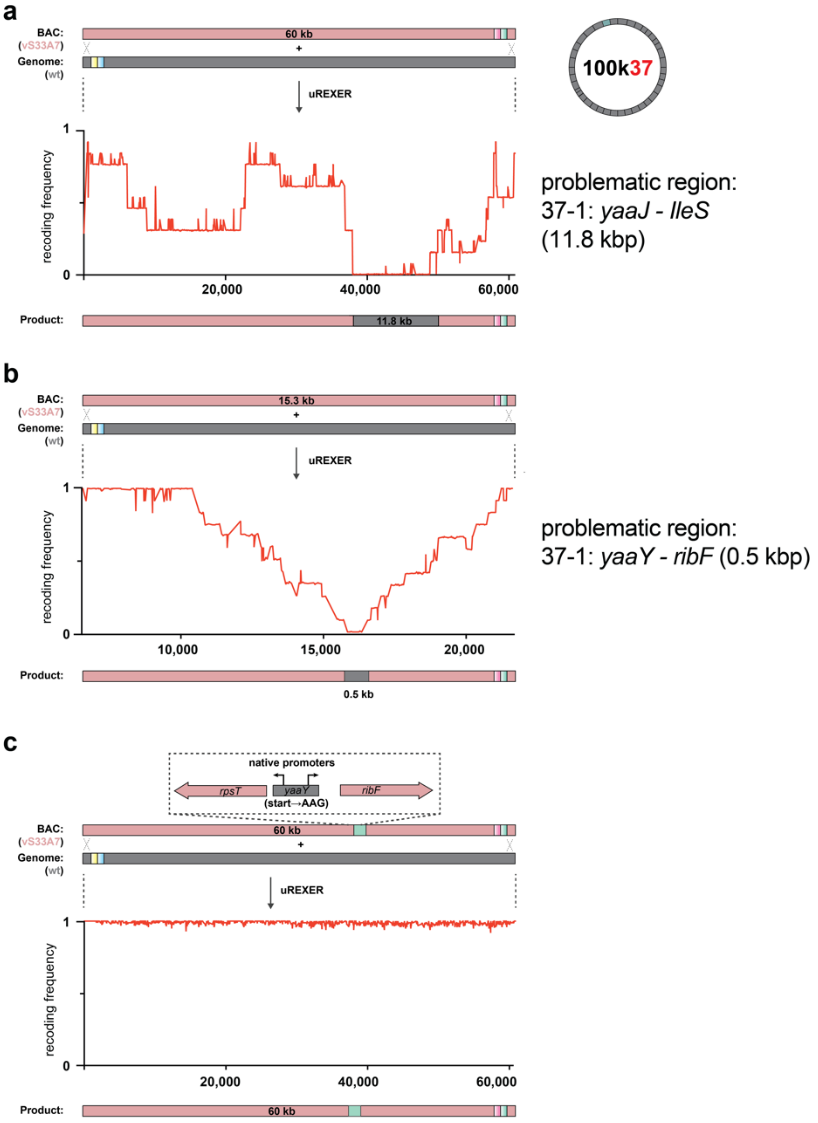


**Supplementary Fig. 12. Fully recoding 100k37 in the genome.**
**a.** Compiled recoding landscape (from 13 clones) from the 100k37 uREXER. The synthetic, recoded, DNA from the BAC (pink) was used for uREXER into the wild-type (grey) genome. The resulting compiled recoding landscape revealed an 11.8-kb region of the genome which was not recoded.
**b.** A miniBAC containing the synthetic recoded DNA covering the region of the genome that was not recoded in the initial uREXER was used for a new uREXER. The resulting compiled recoding landscape (from 12 clones) refined the region of the genome that was not recoded to 0.5 kb (grey). This region contained the full *yaaY* gene and the N-terminus of *ribF*.
**c.** We designed the indicated rational fix which was directly incorporated into the full 100k37 BAC. Integrating this BAC design by uREXER into a wt genome yielded several fully recoded clones (shown is the recoding landscape from an individual clone). The uREXER was performed in a strain also carrying the fully recoded fragment 100k04. Only recoding of 100k37 is shown.

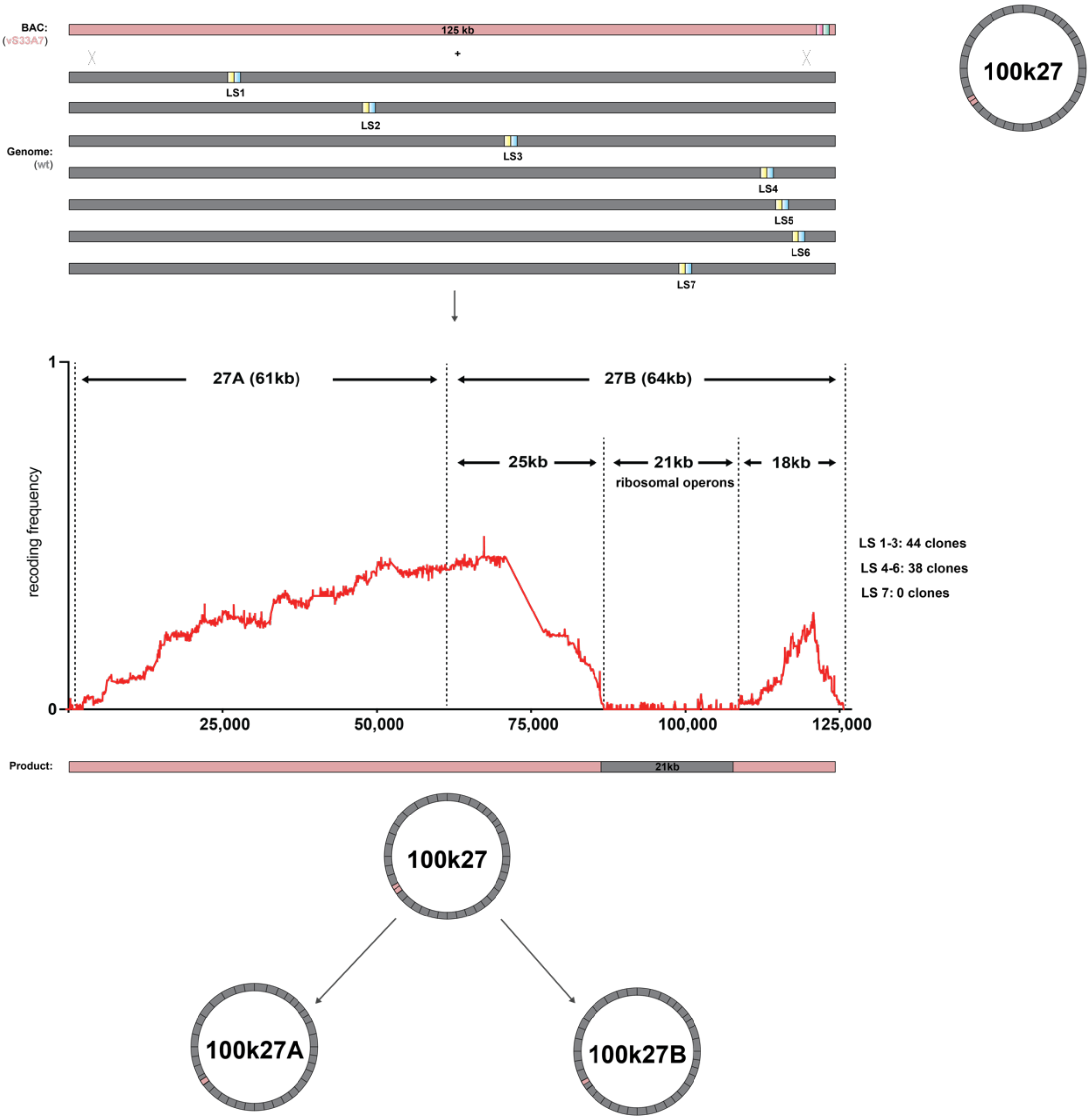


**Supplementary Fig. 13. Mapping the recoding of 100k27 in the genome and defining fragments 27A and 27B.**

Recombination to investigate the recoding of 100k27 in the genome. We integrated *rpsL-KanR* (yellow box, blue box) landing sites at seven separate locations into ΔrecA cells: LS1-3 are located before the ribosomal operons, LS4-6 are located after the ribosomal operons, and LS7 is located within a ribosomal operon upstream of non-essential gene *pioO*. This generated seven landing site strains. We co-transformed the original 100k27 BAC with a helper plasmid carrying arabinose-inducible recA and Cas9 into each landing site strain and used it for recombination. Sequences of correctly phenotyping clones from all seven recombinations (82 clones in total) were used to compile a single recoding landscape. The 21kb region that remains wildtype sequence in the compiled recoding landscape substantially overlaps with the ribosomal protein operons in this region. On the basis of this compiled recoding landscape we divided the sequence into regions 27A (61kb) and 27B (64kb). In subsequent work we recoded 27A and 27B in parallel strains.

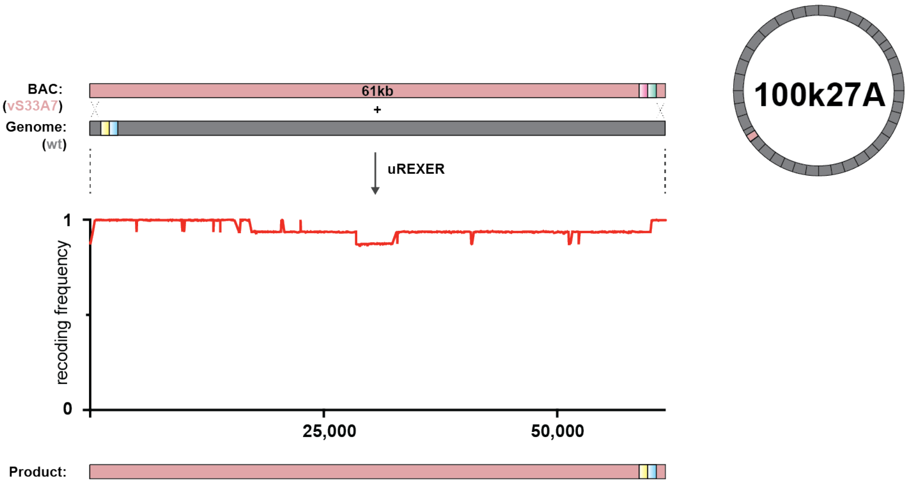


**Supplementary Fig. 14. Fully recoding 61k27A in the genome.**
Compiled recoding landscape (from 14 clones) for the 61kb 100k27A uREXER. The synthetic, recoded, DNA from the BAC (pink) was used for uREXER performed on a clone that already contained genomically recoded 100k26 but wild-type sequence for 100k27 (grey). We identified clones with complete genomic recoding across 27A.

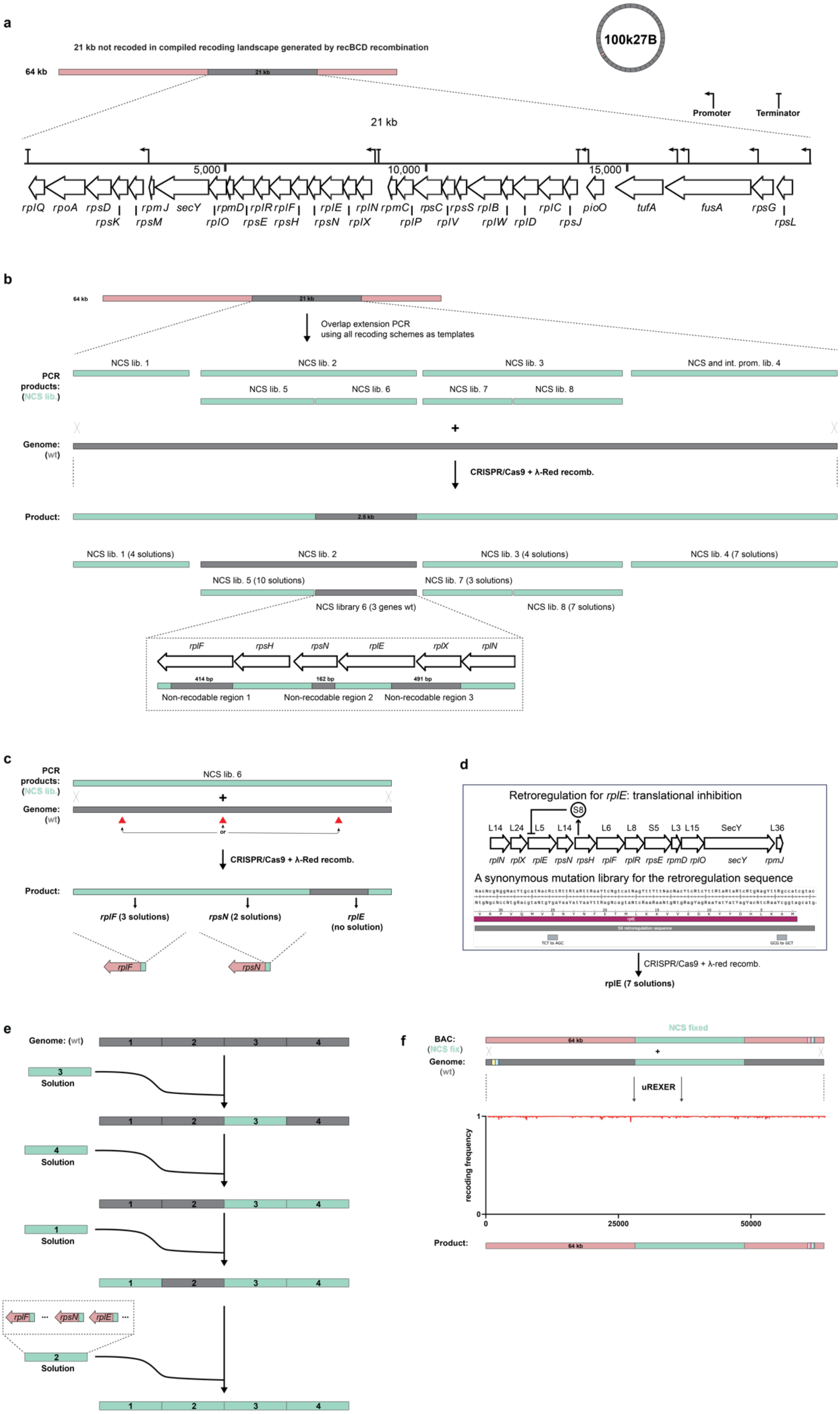
a
21 kb not recoded in compiled recoding landscape generated by recBCD recombination
100k27B
64 kb
21 kb
Promoter
Terminator
5,000
10,000
15,000
rplQ rpoA rpsD rpmJ secY rpmD rplR rplF rplE rplN rpmC rpsC rpsS rplB rplC pioO tufA fusA rpsG
rpsK rpsM rplO rpsE rpsH rpsN rplX rplP rplV rplW rplD rpsJ rpsL
b
Overlap extension PCR using all recoding schemes as templates
NCS lib. 1
NCS lib. 2
NCS lib. 3
NCS and int. prom. lib. 4
PCR products: (NCS lib.)
NCS lib. 5
NCS lib. 6
NCS lib. 7
NCS lib. 8
Genome: (wt)
CRISPR/Cas9 + λ-Red recomb.
Product:
2.5 kb
NCS lib. 1 (4 solutions)
NCS lib. 3 (4 solutions)
NCS lib. 4 (7 solutions)
NCS lib. 5 (10 solutions)
NCS lib. 7 (3 solutions)
NCS library 6 (3 genes wt)
NCS lib. 8 (7 solutions)
rplF rpsH rpsN rplE rplX rplN
414 bp
162 bp
491 bp
Non-recodable region 1
Non-recodable region 2
Non-recodable region 3
c
PCR products: (NCS lib.)
NCS lib. 6
Genome: (wt)
or
CRISPR/Cas9 + λ-Red recomb.
Product:
rplF (3 solutions)
rpsN (2 solutions)
rplE (no solution)
d
Retroregulation for rplE: translational inhibition
S8
L14 L24 L5 L14 L6 L8 S5 L3 L15 SecY L36
rplN rplX rplE rpsN rpsH rplF rplR rpsE rpmD rplO secY rpmJ
A synonymous mutation library for the retroregulation sequence
S8 retroregulation sequence
TCT to AGC
GCG to GCT
CRISPR/Cas9 + λ-red recomb.
rplE (7 solutions)
e
Genome: (wt)
1 2 3 4
Solution
rplF rpsN rplE
f
NCS fixed
BAC: (NCS fix)
Genome: (wt)
uREXER
recoding frequency
0
1
25000
50000
Product:
64 kb

**Supplementary Fig. 15. Fully recoding 64k27B in the genome.**

**a.** 21kb region of the genome (grey) that remained wild-type in the compiled recoding landscape for 100k27 generated by recABCD-mediated recombination. A map of the 21kb region is shown.

**b.** To recode the 21kb region in the genome, we first divided this region into 4 pieces; each piece corresponds to the transcriptional unit from one of the 4 main promoters. We generated PCR products for each piece using four synthetic DNA templates; these templates are the 21-kb region recoded with vS33A7, vS34A7, vS35A7, vS36A7. These PCR products were generated with overlap extension PCR primers for all essential and semi-essential genes (all genes except *rpmJ* and *pioO*) to introduce NCS libraries (and internal promoter libraries of fragment 4). We also split libraries 2 and 3, which contain a large number of genes, into smaller libraries (libraries 5-8) to more effectively cover NCS diversity within genes. The PCR products were used to replace the corresponding genomic sequence by pTarget-based recombination (CRISPR/Cas9 targeting cleavage of each wild-type genomic sequence and lambda-red recombination to integrate the PCR products into the genome). We obtained genomic recoding solutions for all libraries except libraries 2 and 6. Following these experiments only 3 genes (*rplF*, *rpsN, rplE)* within NCS library 6 were never recoded in the genome.

**c.** The diagram shows the gene structure and regions that were not recoded (grey) within the region of the genome covered by NCS library 6 in the experiments described in panel **b**. To obtain solutions for the 3 genes (*rplF*, *rpsN, rplE)* that were not fully recoded, we used NCS libraries targeting each of them. We found solutions for *rplF* and *rpsN*, but not for *rplE*. The experiment was performed by pTarget-based recombination and repeated three times.

**d.** There is a retroregulation sequence at the N-terminus of *rplE*. Ribosomal protein S8 encoded by *rpSH* acts as a repressor for the *rplE* operon. The binding site of S8 spans from -12 to 93 of the *rplE* gene. We therefore designed a larger NCS library and introduced this library into the genome by pTarget-based recombination. From this library we discovered 7 solutions, consistent with code compression, for recoding.

**e.** The solutions for each of the four pieces were introduced into the genome sequentially by pTarget-based recombination. This led to genomic recoding of the 21kb region, with a 73 bp wild-type sequence at the C-terminus of *rplP*; this wild-type sequence was replaced with synthetic sequence that implemented the original genome design by lambda-red recombination.

**f.** Recoding of 64k27B in the genome. We constructed a BAC containing 64 kb of synthetic DNA (64k27B). In this BAC the 21kb solution identified in **e** was flanked by synthetic DNA that implements the original genome design. uREXER of this BAC into the strain generated in **f** led to complete recoding of the entre 64 kb. The recoding landscape from a uREXER clone shows full recoding across the entire fragment.

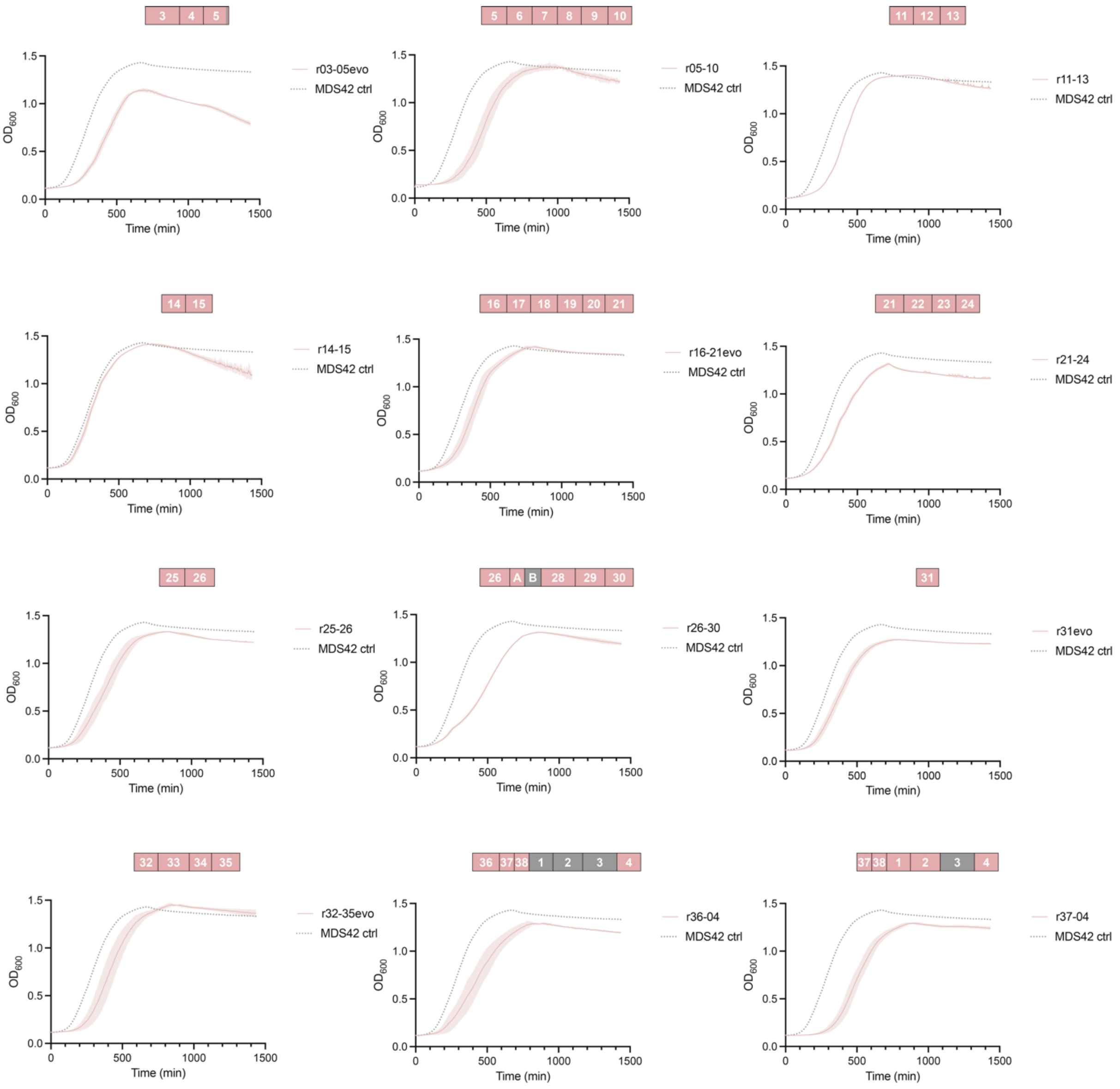


**Supplementary Fig. 16. Growth dynamics of large, recoded sections generated by GENESIS and of evolved strains.**

The growth curves (pink) of the large, recoded sections shown in **Fig. 3** are compared to the growth of the parental strain (MDS42, grey dotted line, the growth curve of parental strain was measured in the same experiment and the same data is plotted as a comparison for every recoded section). The addition of 'evo' to the end of a strain name indicates that the strain contains mutations acquired in an evolution experiment, and apostrophes in the pink boxes indicate the respective fragment of the mutations within the recoded section. Grey boxes indicate non-recoded fragments between recoded fragments. The strains exhibit the following doubling times: r03-05evo: 91 min, r05-10: 104 min, r11-13: 90 min, r14-15: 66 min, r16-21evo: 88 min, r21-24: 96 min, r25-26: 75 min, r26-30: 95 min, r31evo: 75 min, r32-35evo: 84 min, r36-04: 82 min, r37-04: 85 min.

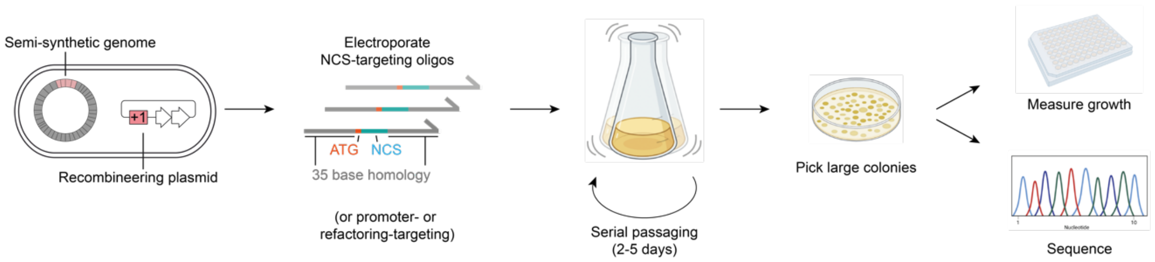


**Supplementary Fig. 17. Overview of the targeted adaptive evolution strategy.**
To improve strain fitness via the targeted mutagenesis of candidate genes that may contribute to reduced fitness, we utilized oligo recombineering together with growth-based selections. Strains harbouring a genome containing recoded sections and a plasmid to enable oligos to recombine into the genome via lagging strand annealing were electroporated with oligo libraries that targeted genes (e.g. the gene N-terminal coding sequences) which were suspected to be associated with a fitness defect when recoded. Subsequently, the cells were passaged to enrich faster growing clones, and plated out to select for large colonies. The growth of individual clones was measured to verify an improvement in fitness, and clones were sequenced to identify which targeted mutations had accumulated over the course of the evolution.

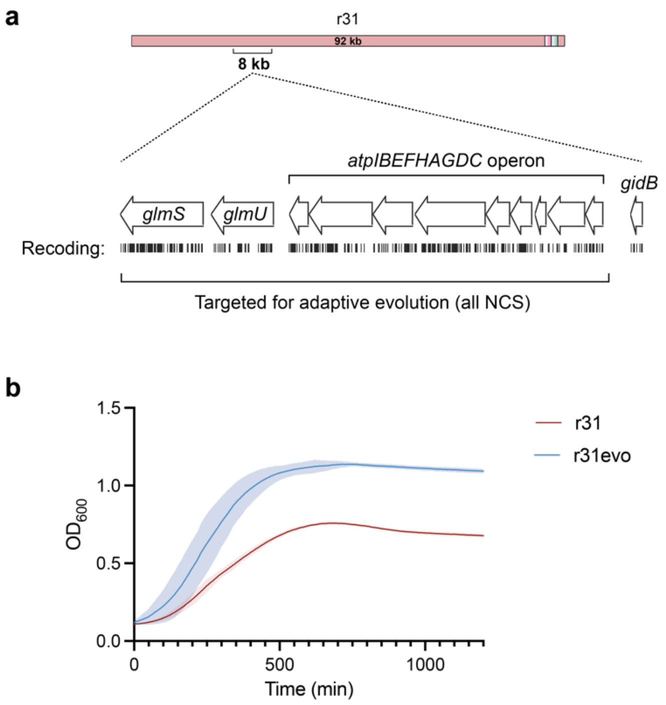


**Supplementary Fig. 18. Improving the growth of genomic 100k31 and generation of r31evo.**

**a.** An 8-kb region in genomically recoded 100k31 with the *atp* and *glm* operons and encompassing the 4.2-kb region associated with slow growth, (**Supplementary Fig. 10**) was targeted for mutation. We targeted the codons for the N-terminal amino acids of all genes in these operons with NCS oligo libraries.

**b.** Growth measurements of the evolved clone taken forward for genome assembly and the strain from which it was derived.

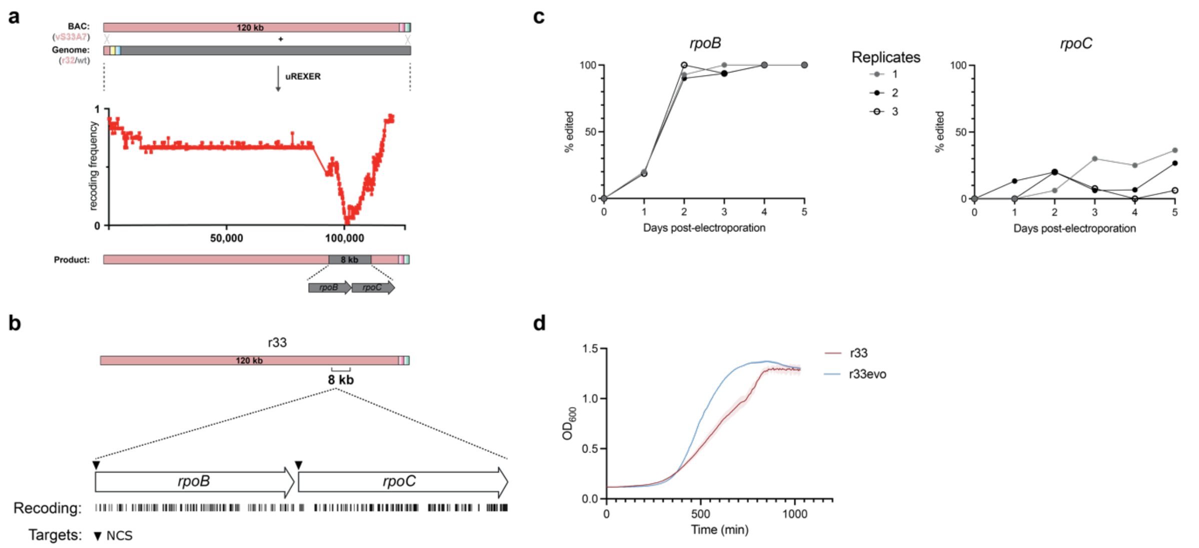


**Supplementary Fig. 19. Targeted adaptive evolution of genomic 100k33 generates 100k33evo.**

**a.** Compiled recoding landscape (from 16 clones) from the 100k33 uREXER. The synthetic, recoded, DNA from the BAC (pink) was used for uREXER into a genome in which 100k32 was recoded. The resulting compiled recoding landscape revealed an 8-kb region with 100k33 which was not recoded – this region contains the *rpoB* and *rpoC* genes.

**b.** Following uREXER-based integrations of recoded 100k33 into a genome in which 100k32 was recoded, we found that the *rpoB* and *rpoC* genes, within 100k33, were not recoded. 100k33 could be fully recoded, however, when performing uREXER in an otherwise non-recoded genome (**Figure 2**). Thus, we sought to evolve the r33 strain, harbouring genomically recoded 100k33, for improved fitness. We targeted both the *rpoB* and *rpoC* genes with oligo-based NCS libraries.

**c.** Assessment of NCS edit accumulation in the *rpoB* and *rpoC* genes. Edit accumulation was assessed by plating out the cultures, picking the largest 48 colonies, and sequencing the genes via Sanger sequencing. Cultures from three electroporations were passaged independently. Edits in the *rpoB* gene enriched to essentially quantitative levels after just two days of passaging.

**d.** Growth measurements of the evolved clone r33evo, containing the sequence of *rpoB* used for further work. The growth of the parental strain, r33 is shown for comparison. The NCS edits selected here were subsequently introduced on the 100k33 BAC via retron-based recombineering, creating 100k33evo BAC; this was used to enable the synthesis of a strain in which 100k32-35 was recoded.

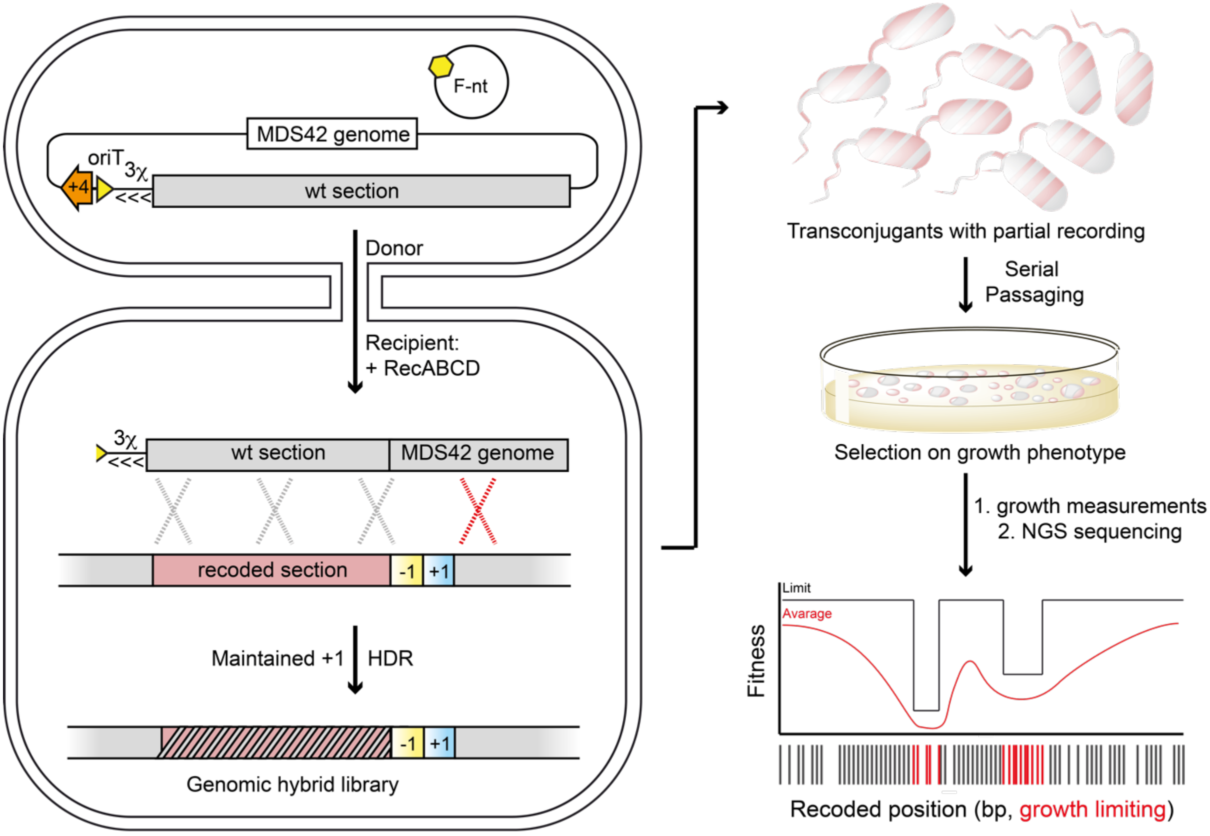


**Supplementary Fig. 20. Linkage Mapping of large, recoded sections.**
To generate a recoding-fitness linkage map of a given recoded genomic section, a donor strain containing the corresponding wild-type section of the parental genome was prepared. In the wt donor strain, a GenR-oriT cassette (GenR: orange, +4, oriT: yellow triangle) was inserted into the genome upstream of the section of interest; this strain also contained a non-transferrable F-plasmid (RK24-oriT::bsd), which mobilizes the genome (from OriT) into the recipient but does not self mobilize. The wt section of the genome was transferred from the donor cell into the recipient cell. Once in the recipient cell, the transferred wt DNA undergoes homologous recombination via the RecABCD pathway, leading to a range of chimeras. Recipient cells were selected for maintenance of the genomic marker in the recipient cells. Crossovers between the recoded genome and the wild-type genome are essentially limited to the region between the oriT and the positive marker on the recoded genome. The genomes of the resulting cells contained chimeras between wt and recoded DNA. The resulting cells were subjected to selection for growth by serial passaging. At regular intervals, single colonies were isolated from the passage culture and colonies with distinct growth phenotypes were selected (usually on the basis of colony size). For the selected colonies, growth curves were measured and key growth parameters determined (doubling time and maximal optical density). The cells were then sequenced by NGS to determine the extent of recoding. The sequence and growth data were compiled into a recoding fitness landscape by calculating the average and limit value (minimum or maximum) at each recoded position across all sequenced clones. Per target codon, the graph shows the average growth parameter across all clones where the respective codon is still recoded. Additionally, for each codon the graph shows the limit value of the growth parameter for all clones tested where that codon is recoded (e.g. maximal optical density or fastest doubling time across all clones).

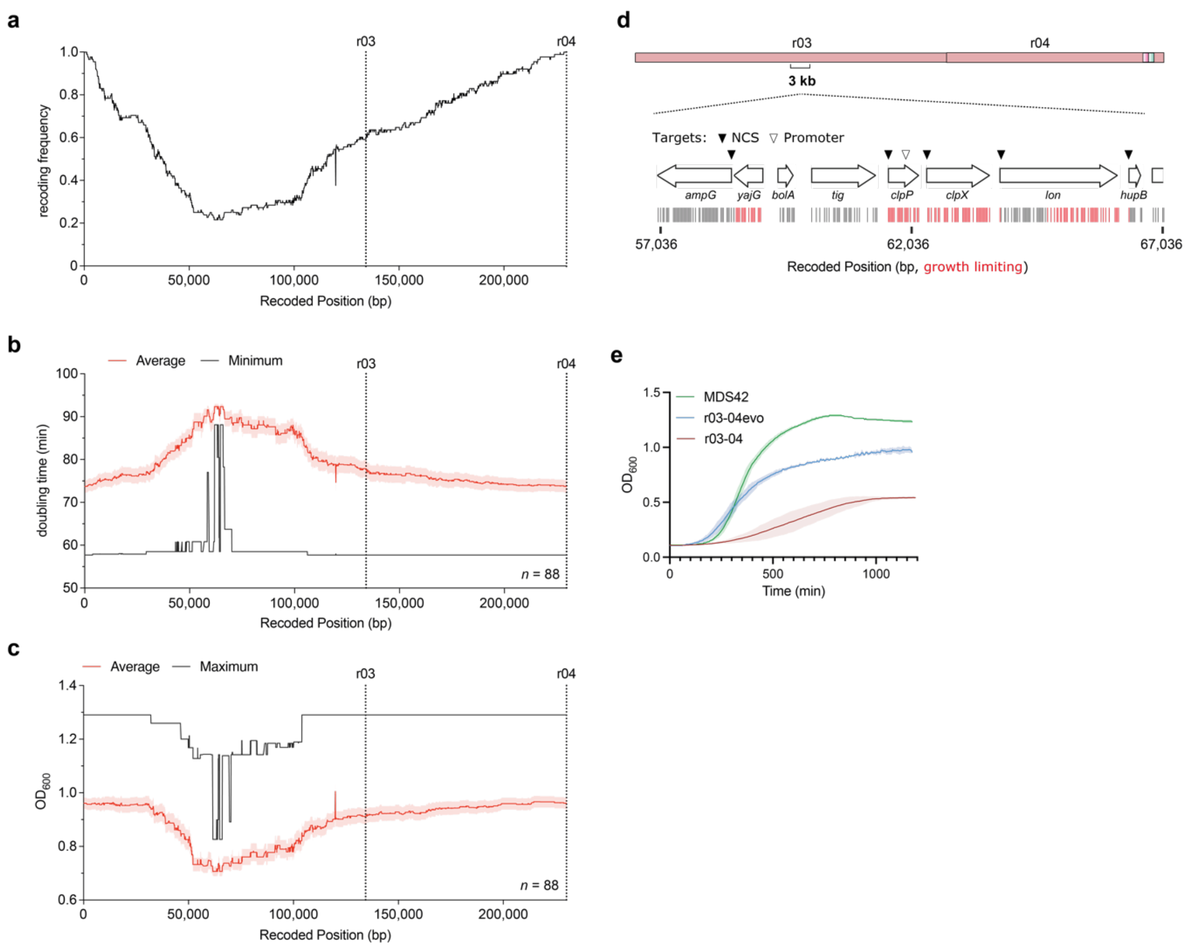


**Supplementary Fig. 21. Linkage mapping and growth improvement for recoded section r03-04.**

**a.** Compiled recoding landscape of all sequenced colonies for the linkage mapping of r03-04. A total of 88 colonies were sequenced. The frequency of recoding at each genomic position is shown.

**b.** Compiled fitness-recoding landscape for doubling time. For each recoded position, the average and limit value (d.t.: minimum) for all sequenced clones is shown. Recoded position numbering is relative to the beginning of the recoded section r03-04 with the distance given in base pairs; dotted lines indicate the end of each individual recoded fragment.

**c.** Compiled fitness-recoding landscape for maximal optical density. For each recoded position the average and limit value for all sequenced clones is shown (max$OD_{600}$: maximum). Recoded position numbering is relative to the beginning of the recoded section r03-04 with the distance given in base pairs; dotted lines indicate the end of each individual recoded fragment.

**d.** We identified a 3-kb region of recoded DNA within r03 that correlates with limited growth. Vertical red lines indicate the recoded codon positions correlated with limit values for d.t. and maximum $OD_{600}$, black lines indicate the positions of other recoded codons in this 3 kb region, the genes are shown as white arrows with their corresponding gene name. Within the mapped region we targeted the N-terminal amino acids of all five genes with NCS oligo libraries and a promoter library targeting the *clpX* promoter within *clpP*.

**e.** Growth comparison of the initial r03-04, the r03-04evo strain taken forward, and the parental strain.

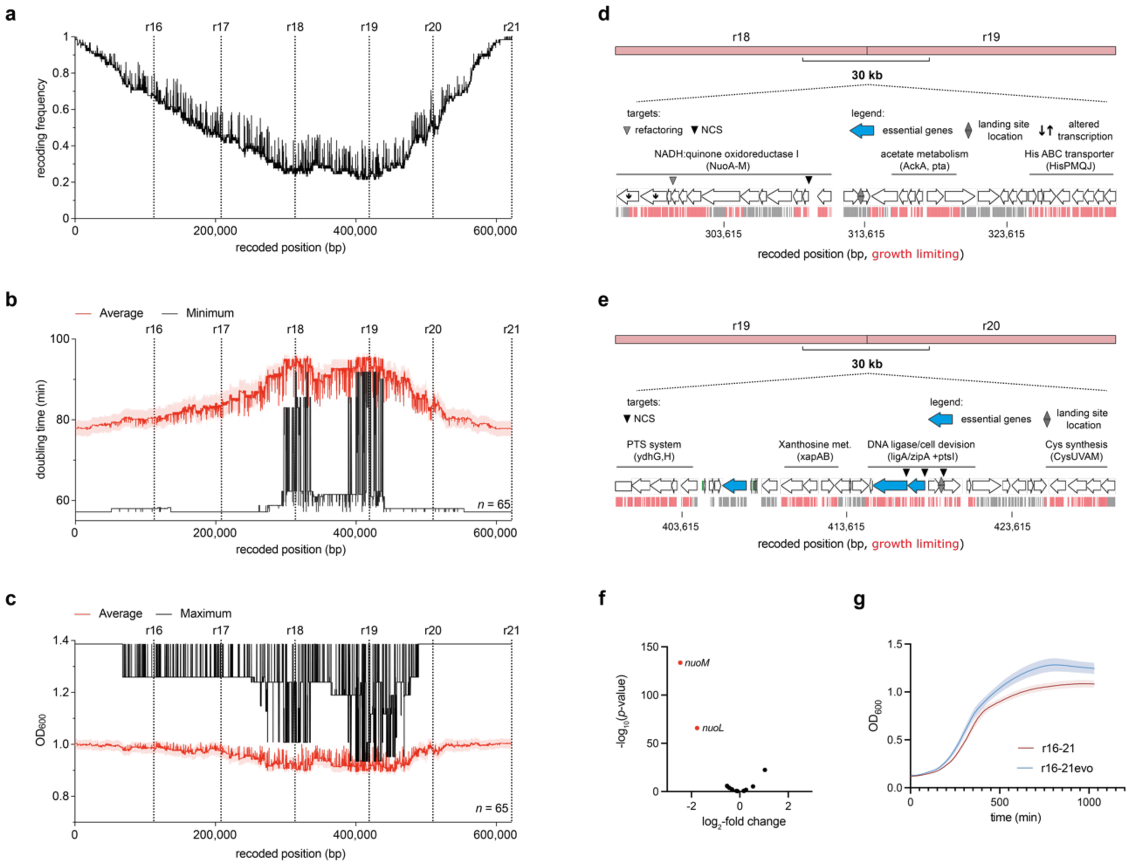

**Supplementary Fig. 22. Linkage mapping and growth improvement fixing of recoded section r16-21.**

**a.** Recoding landscape of all sequenced colonies for the linkage mapping of r16-21. A total of 68 colonies was sequenced. The frequency of recoding at each genomic position is shown.

**b.** Compiled fitness-recoding landscape for doubling time. For each recoded position the average and limit value in all sequenced clones is shown (d.t.: minimum). Recoded position numbering is relative to the beginning of the recoded section r16-21 with the distance given in base pairs; a dotted lines indicate the end of each individual recoded fragment.

**c.** Compiled fitness-recoding landscape for maximal optical density. For each recoded position the average and limit value in all sequenced clones is shown (d.t.: minimum; maxOD$_{600}$: maximum). Recoded position numbering is relative to the beginning of the recoded section r16-21 with the distance given in base pairs; a dotted lines indicate the end of each individual recoded fragment. From the analysis shown in panels **b** and **c** we identified two regions of about 30 kb of recoded DNA at the border between r18 and r19, panel **d**, and r19 and r20, panel **e**, that correlate with limited growth.

**d.** 30-kb region at the border between r18 and r19. Genes are shown as white arrows with operons of interest labelled, recoding events correlated with the limit values for growth are shown as red lines. There are no essential genes (blue) within this mapped region, however the NADH:quinone oxidoreductase I operon is associated with cell growth [63].We decided to target the NCS of the first operon gene *nuoA* and the refactoring event between *nuoJ* and *nuoK*. The refactoring was targeted as the lack of predicted internal ribosomal binding sites and overlapping reading frames indicates operon translation through the RBS-independent termination-reinitiation mechanism (TeRe). This hypothesis was supported by RNA seq data

which identified the reduced transcript levels of genes following the refactoring event (black arrows and panel **f**).
**e.** 30-kb region at the border between r19 and r20. Genes are shown as white arrows with operons of interest labelled, recoding events correlated with the limit values for growth are shown as red lines. We targeted two essential genes (*ligA* and *zipA*) and a regulatory gene (*ptsI*) within the mapped region in close proximity to the previous landing site 19. All genes were targeted with a library of NCS oligos.
**f.** RNA transcript levels measured for the 100k18 BAC relative to the genomic copy. Transcript units are the log2 of their relative fold change, the significance is plotted as the negative log10 of the adjusted p-value for each gene. Only genes within the mapped region in panel **d** are shown. Transcript levels of *nuoM* and *nuoL* were significantly reduced but transcript levels of other genes of the same transcribed unit were unaffected.
**g.** Growth measurements of the evolved clone taken forward for genome assembly and the strain from which it was derived.

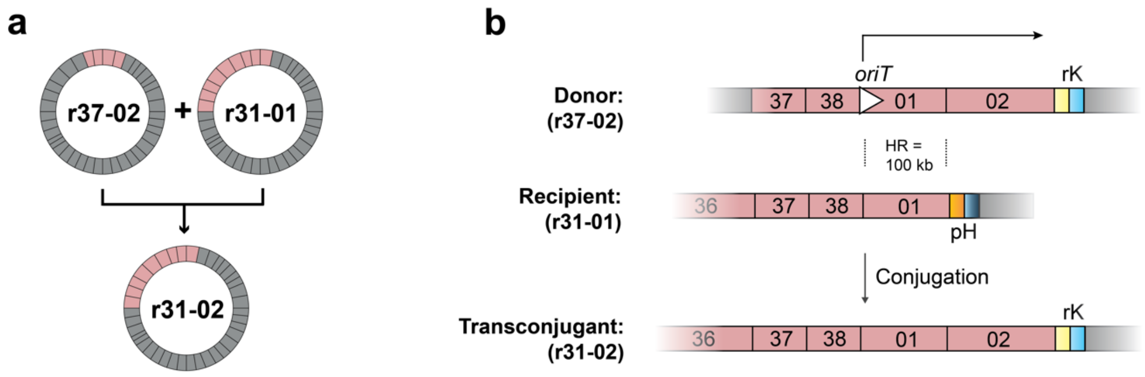


**Supplementary Fig. 23. Conjugation of large recoded sections for the assembly of the recoded genome.**

**a.** Large recoded genomic sections from different strains shown in pink can be combined via conjugation. A conjugation donor (here r37-02) is engineered to conjugate the recoded section into a recipient strain (here r31-01) where the sections recombine to yield a transconjugant strain with a larger recoded genomic section.

**b.** In the donor (r37-02) an origin of transfer (*oriT*) sequence is placed upstream of the section that is to be donated. The donor harbours a positive selection marker at the end of the recoded section and is equipped with a non-transferable F plasmid. The recipient (here r31-01) contains a different double selection cassette (e.g. *pheS-hygR*) that is placed such that it is downstream of a homology region with the donor recoded section (here the homology region (HR) is 100 kb and comprises fragment 100k01). The recipient is additionally equipped with a plasmid containing recA (if not already present on the genome) and a positive selection marker allowing for selection of transconjugants. Conjugation is then initiated and transconjugants can be obtained by selecting for the positive selection marker conjugated by the donor, the positive selection marker harboured on the recipient plasmid, and the negative selection of the outgoing marker downstream of the homology region in the recipient.

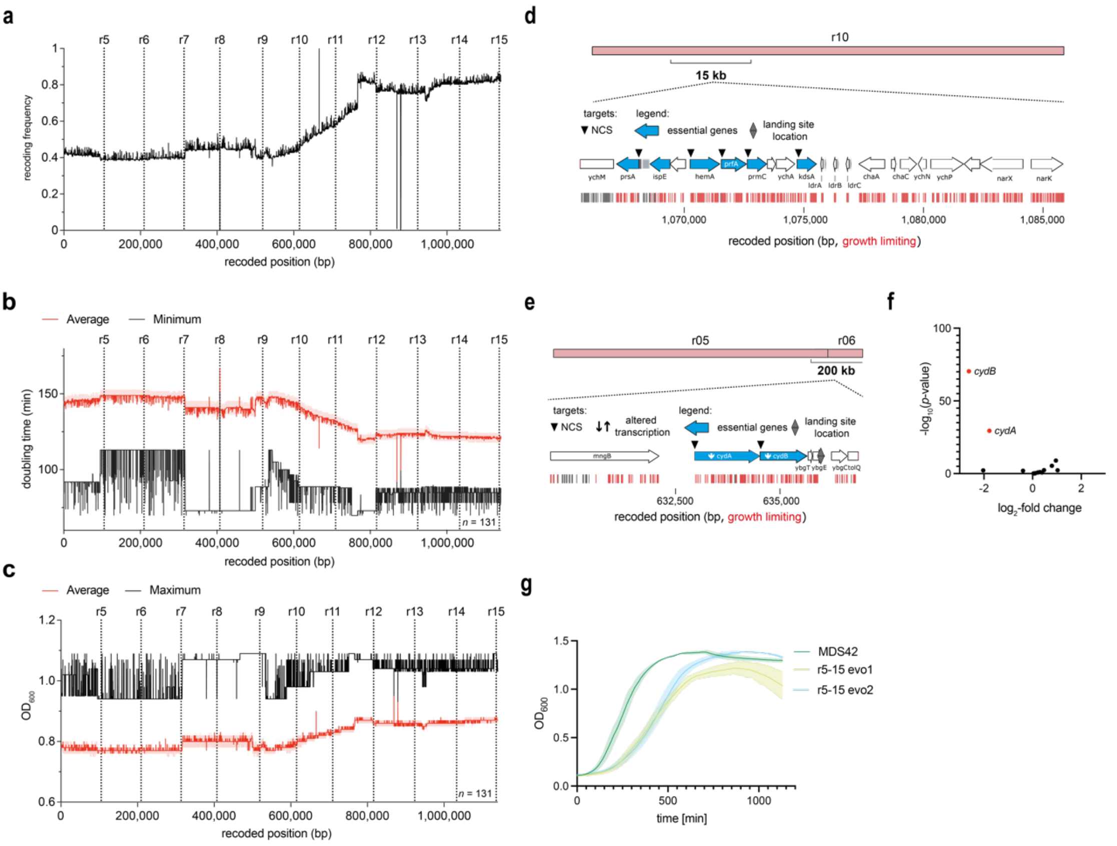


**Supplementary Fig. 24. Recoding-fitness linkage mapping and fixing of recoded section r05-15.**

**a.** Compiled recoding landscape from all 131 sequenced colonies used for the linkage mapping of r05-15. The frequency of recoding at each genomic position is shown.

**b.** Compiled fitness-recoding landscape for doubling time**.** For each recoded position the average and limit value in all sequenced clones is shown (d.t.: minimum); the recoded positions are shown relative to the beginning of the recoded section r05-15 with the distance given in base pairs, dotted lines indicate the end of each individual recoded fragment.

**c.** Compiled fitness-recoding landscape for maximal optical density. For each recoded position the average and limit value in all sequenced clones is shown (max$OD_{600}$: maximum); the recoded positions are shown relative to the beginning of the recoded section 05-15 with the distance given in base pairs, dotted lines indicate the end of each individual recoded fragment.

**d.** From the data in panels **a** and **b** we identified a 15-kb region of recoded DNA within 100k10 of r05-15 that is associated with limited growth. Genes are shown as white arrows with operons of interest labelled, essential genes are highlighted in blue. Recoding events associated with growth limitation are shown as red lines. This 15-kb region was fixed to enable the recoding of 100k10 alone (**Supplementary Fig. 5**). We targeted the N-terminal coding sequences of all essential genes within the mapped region with libraries.

**e.** From the data in panels **a** and **b** we identified the region between the end of 100k05 and 100k07 that is associated with limited growth. We took interest in the indicated 20 kb at the end of 100k05 as it contained two essential genes (blue, *cydAB*) which are in close proximity to the previous landing site location and transcriptionally down regulated. Both genes were targeted with NCS libraries.

**f.** RNA transcript levels measured for the 100k05 BAC relative to the genomic copy. Transcript units are the log2 of their relative fold change, the significance is plotted as the negative log10 of the adjusted p-value for each gene. Only genes within the mapped region in **e** are shown. Recoded *cydA* and *cydB* from the BAC show significantly lower transcription than the non-recoded sequence in the genome.

**g.** Growth measurements of the evolved clone taken forward (r5-15evo2) for genome assembly and the evolution intermediate (r5-15evo1) from which it was derived.

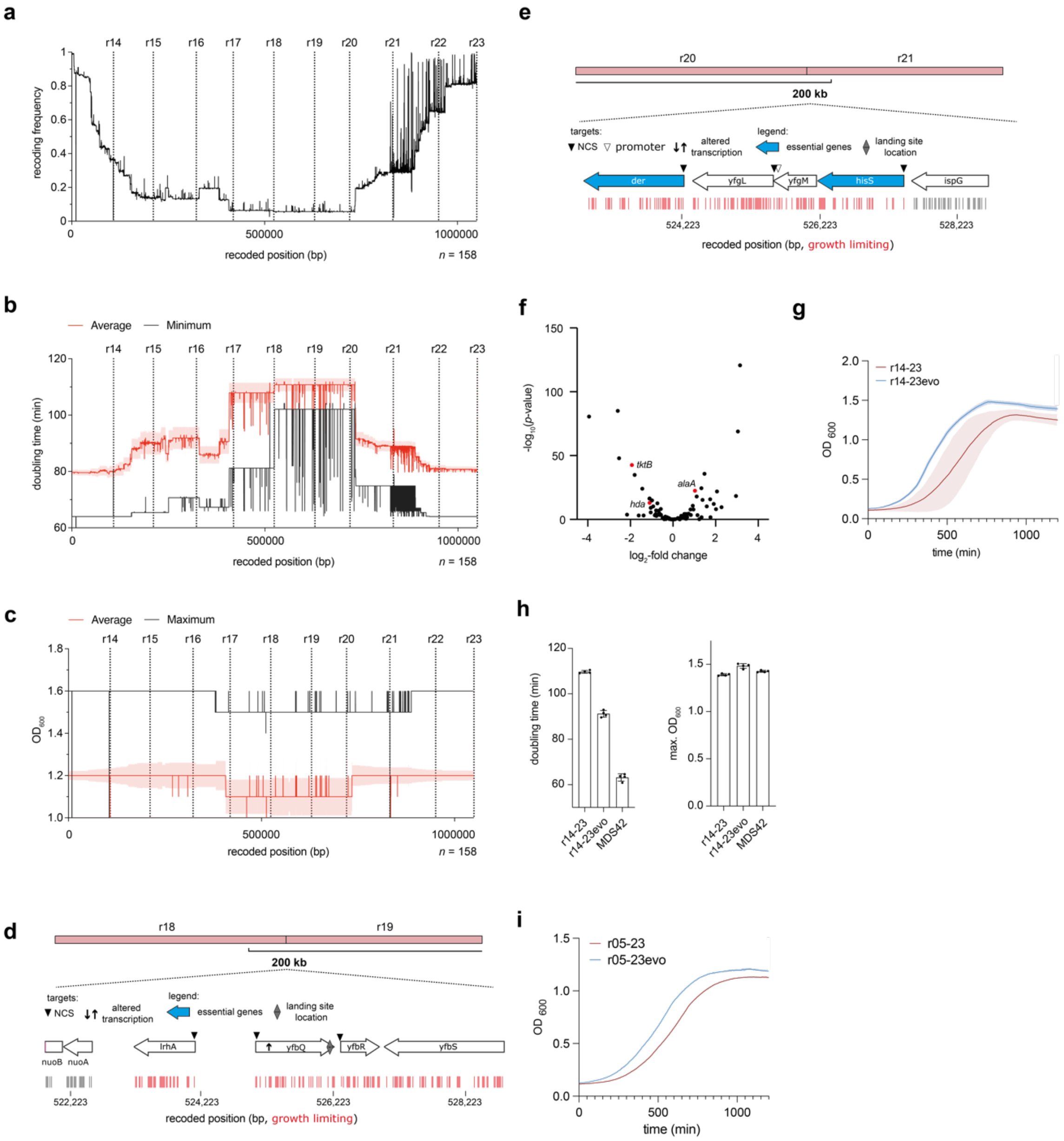


**Supplementary Fig. 25. Recoding-fitness linkage mapping and fixing of recoded section r14-23 and fixing of recoded section r05-23.**

**a.** Compiled recoding landscape from all 158 sequenced colonies used for the linkage mapping of r05-15. The frequency of recoding at each genomic position is shown.

**b.** Compiled fitness-recoding landscape for doubling time**.** For each recoded position the average and limit value in all sequenced clones is shown (d.t.: minimum); the recoded positions are shown relative to the beginning of the recoded section 14-23 with the distance given in base pairs, a dotted lines indicate the end of each individual recoded fragment.

**c.** Compiled fitness-recoding landscape for maximal optical density. For each recoded position the average and limit value in all sequenced clones is shown (d.t.: minimum; maxOD$_{600}$: maximum); the recoded positions are shown relative to the beginning of the recoded section 14-23 with the distance given in base pairs, a dotted lines indicate the end of each individual recoded fragment. These experiments shown in panels **b** and **c** identified a 211.3-kb region between 100k18 and 100k21 where recoding is associated with slow growth.

**d.** Left boundary of region where recoding is associated with slow growth. Genes are shown as white arrows, there are no essential genes in the proximity of the left boundary. Recoding events associated with slow growth are shown as red lines. We decided to target the NCS of *alaA* with a library; the transcript levels for a recoded version of this gene in a BAC were higher than the transcript levels for the non-recoded genomic copy of the gene, shown in panel **f**. Triangles indicate all targeted NCS in proximity of the left boundary.
**e.** Right boundary of region where recoding is associated with slow growth, essential genes are shown in blue. We targeted the operon containing the essential genes *hisS* and *der*, *yfgL* with NCS libraries. Additionally, we targeted the promoter of *yfgL* which was affected by recoding events in *yfgM* with a promoter library. Triangles indicate all targeted NCS and promoters in proximity of the right boundary.
**f.** RNA transcript levels measured for the indicated genes from the BACs containing recoded 100k18, 100k20 and100k21, relative to genomic copy of the genes. Transcript units are the log2 of their relative fold change, the significance is plotted as the negative log10 of the adjusted p-value for each gene. Only genes in the mapped, growth inhibitory region are shown (last gene in 100k18, *alaA*; full 100k20; first 11 kb of 100k21; data for 100k19 were not collected.). Transcript levels of *cysK, purM, purN, tktB, ptsH, ypfE, cchB* and of the essential gene *hda* were significantly reduced (blue).
**g.** Growth measurements of the evolved clone (r14-23evo) and the parental strain from which it was derived.
**h.** Doubling time of the evolved clone and the parental strain from which it was derived. Maximum OD of the evolved clone and the parental strain from which it was derived.
**i.** Growth measurements of the evolved clone r05-23evo and the parental strain from which it was derived. The mapping and evolution on the recoded region 14-23 informed the evolution experiment for the larger recoded region of 05-23 – this yielded the evolved clone r05-23evo taken forward.

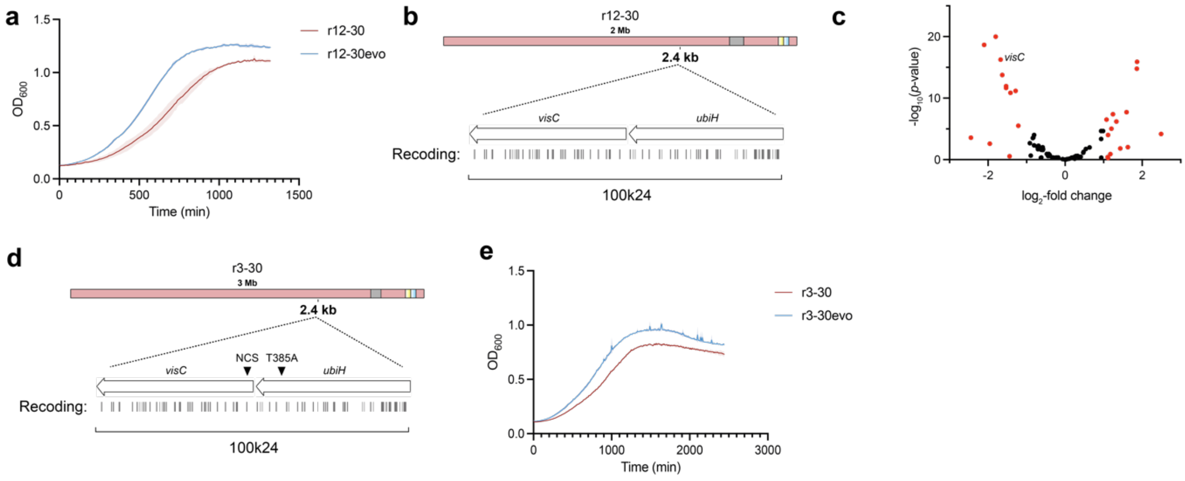


**Supplementary Fig. 26. Evolution of strain r03-30.**
**a.** Passaging of the strain r12-30 resulted in the discovery of an evolved r12-30 strain, which picked up a random mutation in *ubiH*.
**b.** *ubiH* forms an operon with *visC* located in section 24, we hypothesized that the selected point mutation influences expression of *visC*.
**c.** RNA transcript levels measured for the recoded BAC for 100k24 indicated that *visC* is substantially down regulated. Transcript units are the log2 of their relative fold change, the significance is plotted as the negative log10 of the adjusted p-value for each gene.
**d.** We tested whether the random mutation can be recapitulated in r03-30 or whether targeting *visC* directly by an NCS library can improve strain growth.
**e.** Growth measurements of an evolved version (evo) of r03-30 in which the *ubiH* mutation was reintroduced.

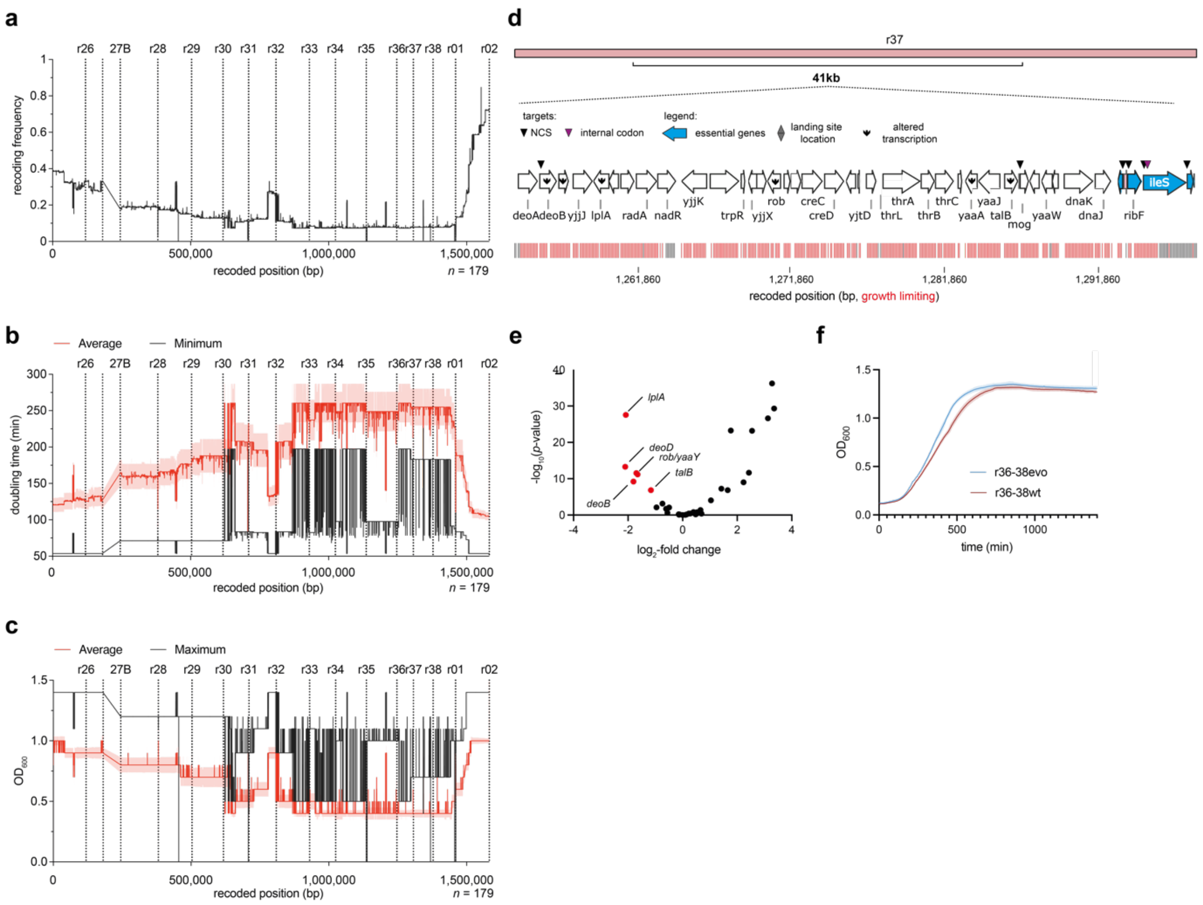


**Supplementary Fig. 27. Recoding-fitness linkage mapping and fixing of recoded section r26-02.**

**a.** Compiled recoding landscape from all 179 sequenced colonies used for the linkage mapping of r26-02. The frequency of recoding at each genomic position is shown. r26-02 is Δ*recA*.

**b.** Compiled fitness-recoding landscape for doubling time. For each recoded position the average and limit value in all sequenced clones is shown (d.t.: minimum); the recoded positions are shown relative to the beginning of the recoded section with the distance given in base pairs, dotted lines indicate the end of each individual recoded fragment.

**c.** Compiled fitness-recoding landscape for maximal optical density. For each recoded position the average and limit value in all sequenced clones is shown (max$OD_{600}$: maximum); the recoded positions are shown relative to the beginning of the recoded section with the distance given in base pairs, dotted lines indicate the end of each individual recoded fragment.

**d.** Through the linkage mapping shown in panels **b** and **c** we identified a 41-kb region of recoded DNA within 100k37 of r26-02 that is associated with limited growth. Genes are shown as white arrows with operons of interest labelled, essential genes are highlighted in blue. Recoding events associated with growth limitation are shown as red lines. We targeted the coding sequences of *deoB*, *mog*, *rpsT*, *ribF*, and *ileS* labelled with a triangle within the mapped region for mutagenesis.

**e.** RNA transcript levels measured for the 100k37 BAC relative to the genomic copy. Transcript units are the log2 of their relative fold change, the significance is plotted as the negative log10 of the adjusted p-value for each gene. Only genes within the 41 kb mapped

region in **d** are shown. Recoded *deoB* from the BAC shows significantly lower transcription than the non-recoded sequence in the genome.
**f.** Growth measurements of the evolved clone taken forward for genome assembly and the parental strain from which it was derived.

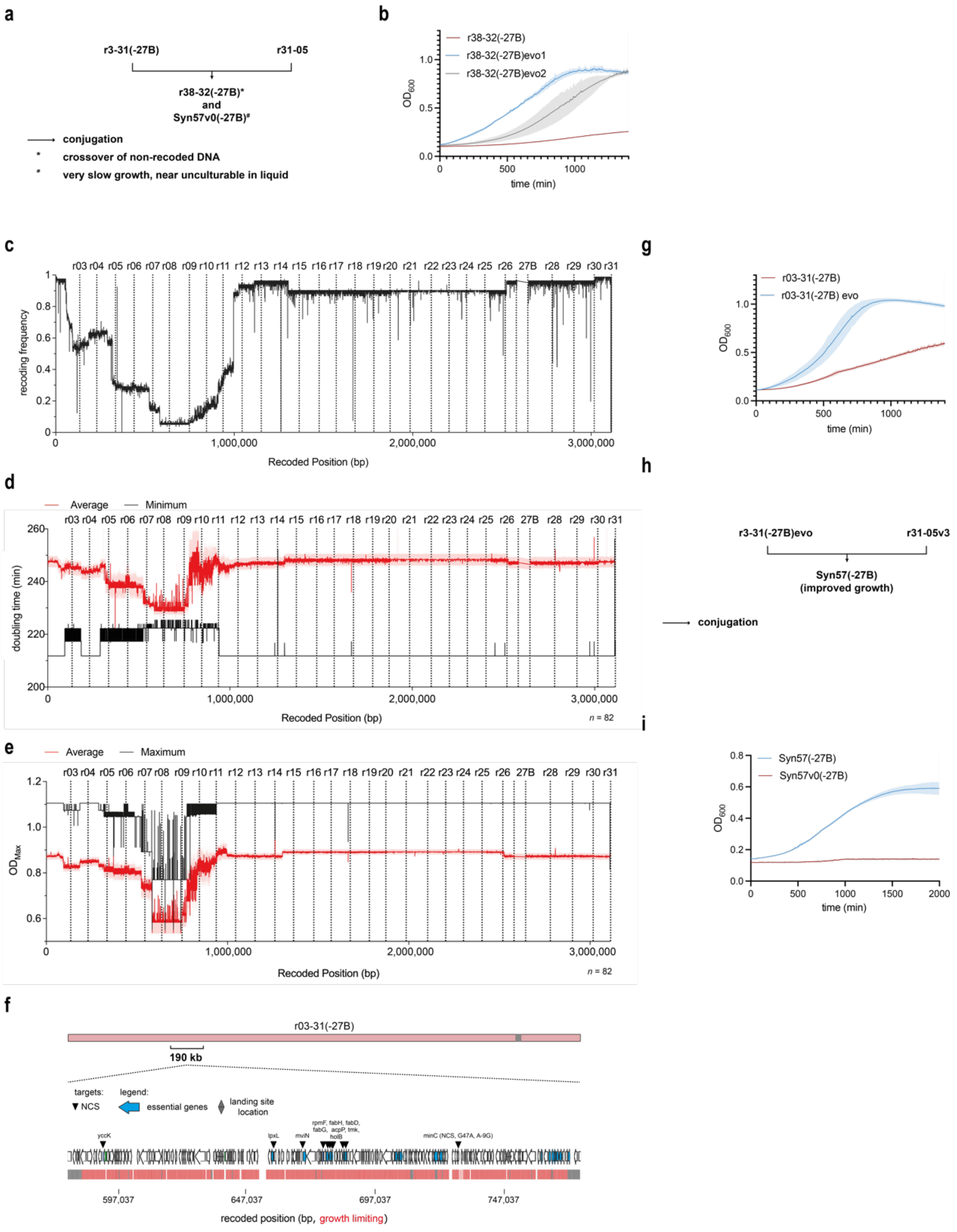

**Supplementary Fig. 28. Linkage mapping and growth improvement fixing of recoded section r03-31.**

**a.** Sections r03-31(-27B) and r31-05 were combined via a conjugative assembly step to yield r38-32(-27B) and Syn57v0(-27B).

**b.** Passaging of the r38-32 strain yielded two clones with improved growth that had enriched different random mutations in *minC*.

**c.** Compiled recoding landscape from all 82 sequenced colonies used for the linkage mapping of r03-31. The frequency of recoding at each genomic position is shown.

**d.** Compiled fitness-recoding landscape for doubling time**.** For each recoded position the average and limit value in all sequenced clones is shown (d.t.: minimum); the recoded positions are shown relative to the beginning of the recoded section with the distance given in base pairs, dotted lines indicate the end of each individual recoded fragment.
**e.** Compiled fitness-recoding landscape for maximal optical density. For each recoded position the average and limit value in all sequenced clones is shown (maxOD$_{600}$: maximum); the recoded positions are shown relative to the beginning of the recoded section with the distance given in base pairs, dotted lines indicate the end of each individual recoded fragment.
**f.** From the analysis shown in panels **d** and **e** we identified a 190 kb region of recoded DNA across r08 to r10 that was linked to a decrease in maximum optical density. We targeted the N-terminal coding sequences of the genes indicated. We also targeted the introduction of the same mutations which we identified in the r38-32 evolution.
**g.** An evolved strain (r03-31(-27B)evo) with substantially improved growth compared to the parent strain was identified.
**h.** The evolved r03-31(-27B) strain was used in a conjugative assembly with r31-05v3 (**Fig. 4**) to yield Syn57(-27B).
**i.** Growth measurements of the new Syn57(-27B) strain compared to the initial Syn57v0 (-27B) strain.

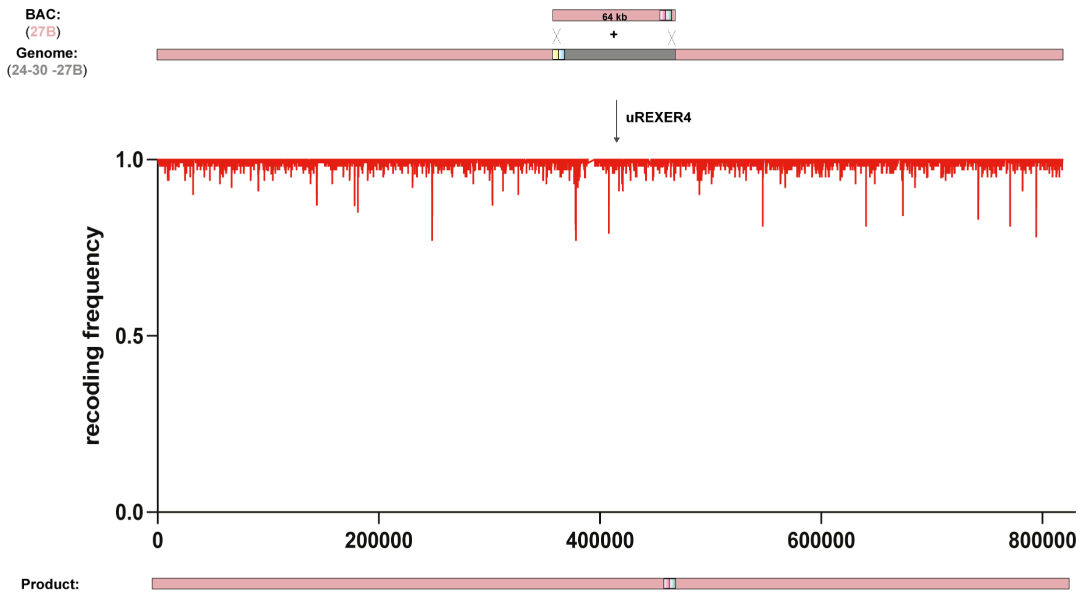


**Supplementary Fig. 29. Genomic integration of 100k27B into the recoded r24-30(-27B) strain.**

After 100k27B was integrated into the MDS42 strain, passaging of the strain accumulated three point mutations in *fusA* and *rpsG* which are compatible with our recoding scheme. A new BAC was constructed, including these mutations. The same sacB-Cm selection markers were used for the BAC, and we integrated a pheS-HygR landing site at the end of 27A in the r24-30(-27B) partially recoded strain. A uREXER4 experiment, which introduced four Cas9 cleavage sites (two universal spacers to excise the recoded region from the BAC and two custom spacers targeting the regions immediately flanking the integration site on the genome), was performed with the new version of the 27B BAC in the r24-30(-27B) strain. One clone was fully recoded, resulting in the complete r24-30 strain.

**Supplementary Data 1 (separate file)**
Recoding schemes for the 20 kb REXERs.
